# Global Absence and Targeting of Protective Immune States in Severe COVID-19

**DOI:** 10.1101/2020.10.28.359935

**Authors:** Alexis J. Combes, Tristan Courau, Nicholas F. Kuhn, Kenneth H. Hu, Arja Ray, William S. Chen, Simon J. Cleary, Nayvin W. Chew, Divyashree Kushnoor, Gabriella C. Reeder, Alan Shen, Jessica Tsui, Kamir J. Hiam-Galvez, Priscila Muñoz-Sandoval, Wandi S Zhu, David S. Lee, Yang Sun, Ran You, Mélia Magnen, Lauren Rodriguez, Aleksandra Leligdowicz, Colin R. Zamecnik, Rita P. Loudermilk, Michael R. Wilson, Chun J. Ye, Gabriela K. Fragiadakis, Mark R. Looney, Vincent Chan, Alyssa Ward, Sidney Carrillo, The UCSF COMET Consortium, Michael Matthay, David J. Erle, Prescott G. Woodruff, Charles Langelier, Kirsten Kangelaris, Carolyn M. Hendrickson, Carolyn Calfee, Arjun Arkal Rao, Matthew F. Krummel

**Affiliations:** Department of Pathology, San Francisco, 513 Parnassus Ave, HSW512, San Francisco, CA 94143-0511, USA; ImmunoX Initiative, San Francisco, 513 Parnassus Ave, HSW512, San Francisco, CA 94143-0511, USA; UCSF CoLabs, San Francisco, 513 Parnassus Ave, HSW512, San Francisco, CA 94143-0511, USA; Department of Radiation Oncology, San Francisco, 513 Parnassus Ave, HSW512, San Francisco, CA 94143-0511, USA; Division of Pulmonary and Critical Care Medicine, Department of Medicine and the Cardiovascular Research Institute, San Francisco, 513 Parnassus Ave, HSW512, San Francisco, CA 94143-0511, USA; Departments of Otolaryngology, San Francisco, 513 Parnassus Ave, HSW512, San Francisco, CA 94143-0511, USA; Department of Microbiology & Immunology, San Francisco, 513 Parnassus Ave, HSW512, San Francisco, CA 94143-0511, USA; Sandler Asthma Basic Research Center, San Francisco, 513 Parnassus Ave, HSW512, San Francisco, CA 94143-0511, USA; Institute of Human Genetics and Division of Rheumatology, Department of Medicine, San Francisco, 513 Parnassus Ave, HSW512, San Francisco, CA 94143-0511, USA; Weill Institute for Neurosciences, Department of Neurology, San Francisco, 513 Parnassus Ave, HSW512, San Francisco, CA 94143-0511, USA; Division of Infectious Disease, Department of Medicine, San Francisco, 513 Parnassus Ave, HSW512, San Francisco, CA 94143-0511, USA; Division of Hospital Medicine, San Francisco, 513 Parnassus Ave, HSW512, San Francisco, CA 94143-0511, USA

## Abstract

While SARS-CoV-2 infection has pleiotropic and systemic effects in some patients, many others experience milder symptoms. We sought a holistic understanding of the severe/mild distinction in COVID-19 pathology, and its origins. We performed a whole-blood preserving single-cell analysis protocol to integrate contributions from all major cell types including neutrophils, monocytes, platelets, lymphocytes and the contents of serum. Patients with mild COVID-19 disease display a coordinated pattern of interferon-stimulated gene (ISG) expression across every cell population and these cells are systemically absent in patients with severe disease. Severe COVID-19 patients also paradoxically produce very high anti-SARS-CoV-2 antibody titers and have lower viral load as compared to mild disease. Examination of the serum from severe patients demonstrates that they uniquely produce antibodies with multiple patterns of specificity against interferon-stimulated cells and that those antibodies functionally block the production of the mild disease-associated ISG-expressing cells. Overzealous and auto-directed antibody responses pit the immune system against itself in many COVID-19 patients and this defines targets for immunotherapies to allow immune systems to provide viral defense.

**One Sentence Summary:** In severe COVID-19 patients, the immune system fails to generate cells that define mild disease; antibodies in their serum actively prevents the successful production of those cells.

---

To understand immune biology amongst COVID-19 patients, we compared them to patients presenting with similar respiratory symptoms but who were not infected with the SARS-CoV-2 virus. We prospectively enrolled 21 SARS-CoV-2 positive inpatients, 11 inpatients with similar clinical presentations consistent with acute lung injury (ALI) or acute respiratory distress syndrome (ARDS), who were SARS-CoV-2 negative—those caused by other infections or of unknown origin—and 14 control individuals. We further categorized these over the next weeks as ‘mild/moderate’ (M/M: typically short stays in hospital with no need for mechanical ventilation and intensive care) or ‘severe’ (requiring intubation and intensive care) according to the full clinical course of their disease (**Fig 1A/S1A** and **Table S1**). Hence, our study includes patients with mild/moderate (n=11) or severe (n=10) COVID-19 and patients with mild/moderate (n=6) or severe (n=5) non-COVID-19 ALI/ARDS. With the exception of one individual, all our patients who presented with mild/moderate disease remained mild/moderate during hospitalization (**Fig S1A**), suggesting that mild/moderate and severe are more stable states rather than transient phases of disease in this cohort.

**Figure 1:**
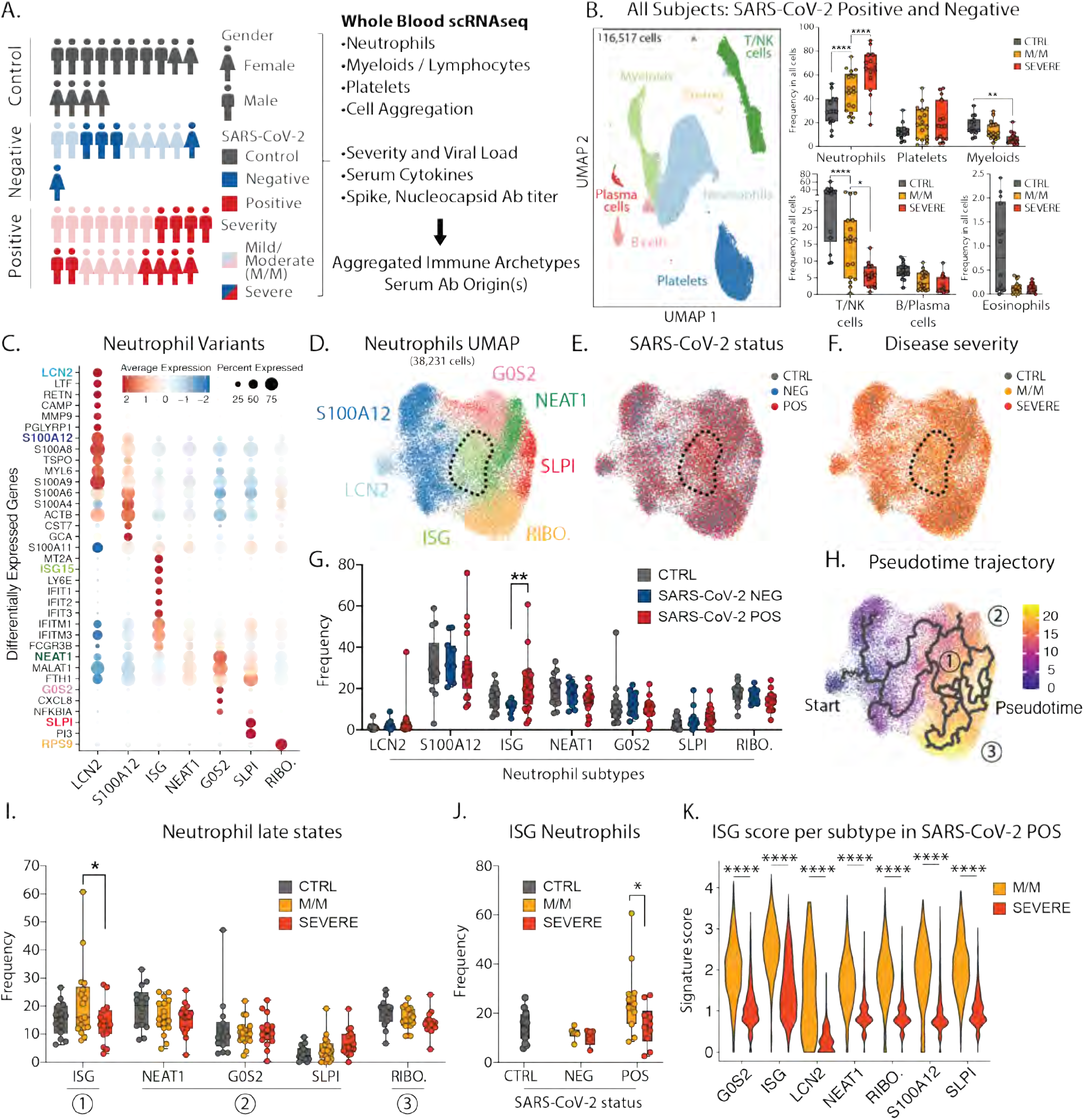
Severe COVID-19 disease is characterized by the lack of IFN-responsive neutrophils. **A**. Gender, SARS-CoV-2 status and disease severity in patients and control individuals (left) and description of study design (right). **B**. UMAP visualization of cells merged from the entire cohort with specific populations overlaid (left), and frequencies of these populations across control, mild/moderate (M/M) and severe individuals (right). **C**. Dotplot representation of top differentially-expressed-genes (DEG) between neutrophil subsets. **D**. UMAP visualization of neutrophil subsets. **E. and F**. Overlay of SARS-CoV-2 status and disease severity, respectively, on the neutrophil UMAP. **G**. Frequencies of neutrophil subsets among all neutrophils across control, SARS-CoV-2 negative and SARS-CoV-2 positive individuals. **H**. Pseudotime trajectory of neutrophil subsets. **I**. Frequencies of the neutrophil subsets among all neutrophils at later stages of pseudotime trajectories across control, mild/moderate and severe individuals. **J**. Frequency of ISG neutrophils among all neutrophils across SARS-CoV-2 status and disease severity. **K.** Score of ISG signature across neutrophil subtypes and disease severity in SARS-CoV-2 positive patients. Statistical significance was assessed using a two-way ANOVA test with multiple comparisons for panels B, G, I and J, and using a Wilcoxon test for panel K. * p-value < 0.05; ** p-value < 0.01; *** p-value < 0.001; **** p-value < 0.0001.

Since the majority of COVID-19 mortality is among patients with the (ARDS)— characterized by an exuberant immune response with prominent contributions from neutrophils, monocytes, platelets—we focused upon faithfully collecting these cells along with other major populations. We thus processed early morning blood samples from all individuals within 3 hours of sampling, and after red blood cell lysis, we analyzed the remaining white blood cells by single-cell RNA sequencing (scRNA-seq). After merging, batch-correction and doublet-removal our data comprised 116,517 cells (**Fig 1B/S1B**) among which we identified neutrophils, platelets, mononuclear phagocytes, T/NK cells, B cells, plasma cells and eosinophils (**Fig S1C**). We confirmed a positive association between neutrophil count and disease severity and an inverse correlation for lymphoid populations (**Fig 1B/S1D**) (*1–3*). At this level of resolution, findings were similar between SARS-CoV-2 negative and positive individuals (**Fig S1E).**

Within the neutrophils, we identified seven subtypes (**Fig 1C/D**), consistent with previous studies (*2, 4*). One population, harboring a strong interferon-stimulated gene (ISG) signature and henceforth termed ISG neutrophils, was highly enriched in SARS-CoV-2 positive patients but not in those whose disease was severe (**Fig 1E/F/G**). Analysis of populations using a pseudotime method to estimate differentiation trajectories (*5*) assigned the starting population as the stem LCN2 population (**Fig 1H/S1F-G**) and suggested three putative late populations: the ISG-expressing population (state 1), a collection of populations sharing expression of NEAT1, MALAT1 and FTH1 (state 2), and a population enriched for ribosomal genes (RIBO.; state 3) which may be en route to cell death. Of these late stages, the ISG subtype was the only one found significantly altered between mild/moderate and severe patients (**Fig 1I**) and specifically within the SARS-CoV-2 positive individuals (**Fig 1J/S1H**). ISG signature genes include master anti-viral regulators such as ISG15 and IFITM3 which restricts viral entry into the cytosol (*6*).

*We* also undertook a second form of analysis of differentially expressed genes (DEG) from SARS-CoV-2 positive versus negative patients, and from mild/moderate versus severe patients across all neutrophils (**Fig S1I-L**). This demonstrated that ISG signature genes are differentially higher in all neutrophils, of all subsets, specifically in SARS-CoV-2 positive mild/moderate patients, than in SARS-CoV-2 positive patients with severe disease (**Fig 1K/S1M-O**). In contrast, a separate neutrophil degranulation gene program is upregulated in mild/moderate as compared to severe disease regardless of COVID status (**Fig S1P-Q**). This suggests a shared program of degranulation enhancement in all respiratory infections regardless of causative pathogen, and a global rise of the ISG program in all neutrophils in mild/moderate SARS-CoV-2 positive cases that is absent in severe SARS-CoV-2 infection (*3*).

Assessing the mononuclear phagocytes—monocytes, macrophages, dendritic cells and plasmacytoid dendritic cells (pDC)—yielded 7 clusters of transcriptionally distinct cells subsets, evenly distributed across our cohort with a heterogenous number of genes and unique molecular identifiers detected for each cluster (**Fig 2A-B/2SA-D**). We identified ISG classical monocytes as being enriched in SARS-CoV-2 positive patients, and particularly those having mild/moderate disease, similarly to neutrophils (**Fig 2B-C/S2E-G**). pDCs which are substantial producers of the cytokine IFNα are also typically elevated in mild/moderate SARS-CoV-2 positive patients although this falls short of statistical significance in our dataset (**Fig 2C**). In contrast, elevated DCs that were previously considered a hallmark of COVID-19 patients when compared to healthy controls (*2*) are also elevated in SARS-CoV-2 negative patients (**Fig 2C**). ISG monocytes also expressed genes associated with glycolysis, compared to the S100A12 subset that were enriched for genes associated with OxPhos metabolism, consistent with previous reports in bacterial sepsis (*7*) (**Fig S3A**). As for neutrophils, differential gene expression analysis demonstrated that ISGs were the dominant genes associated with mild/moderate phenotypes when the entire mononuclear phagocyte pool was assessed (**Fig 2D**).

**Figure 2:**
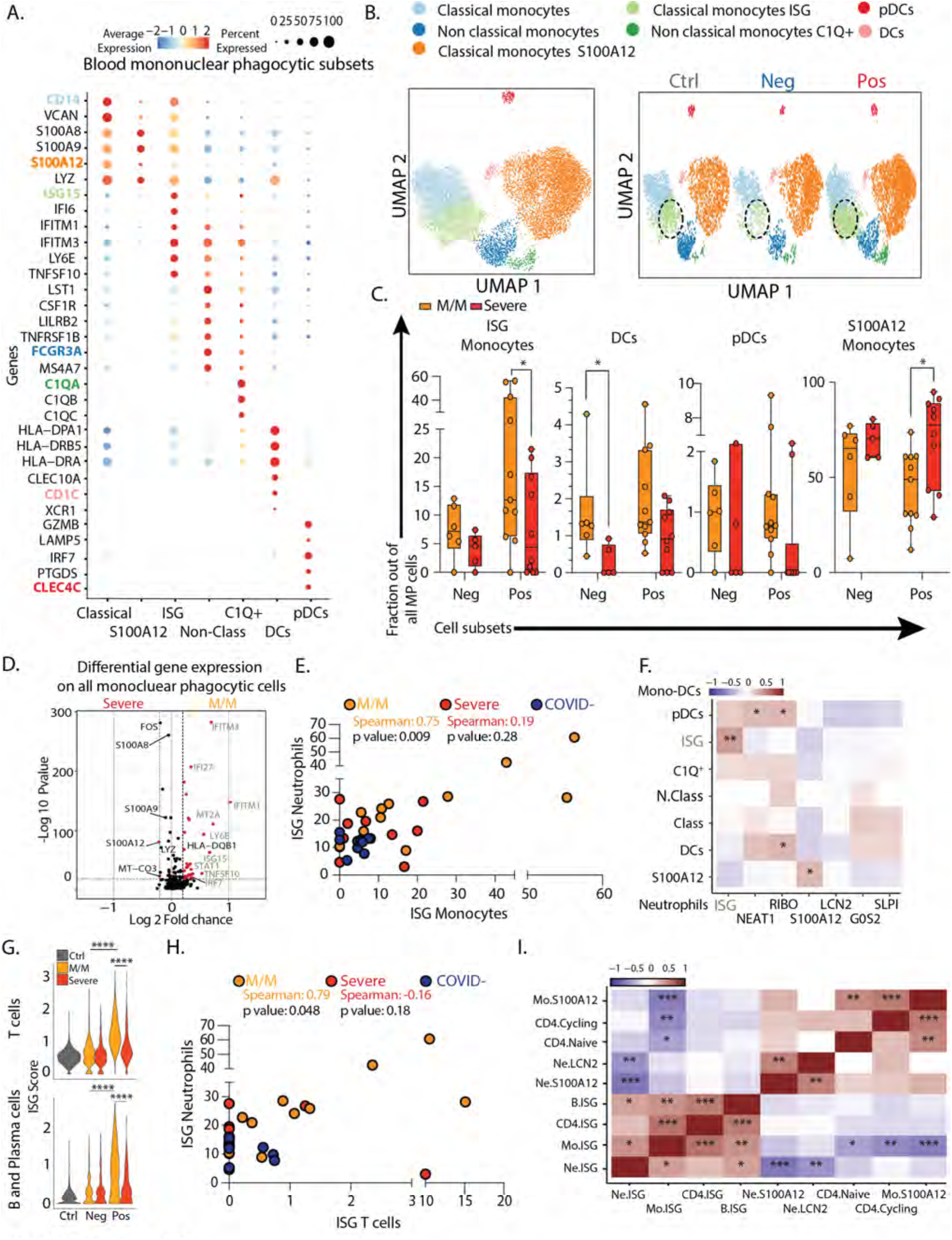
Severe COVID-19 disease is defined by the lack of a concerted IFN-response across peripheral blood immune cells. **A**. Dotplot representation of the top differentially-expressed-genes (DEG) between clusters identified in blood mononuclear phagocytic cell (MPC) subsets. **B**. UMAP visualization of the 19,289 MPC isolated from the entire dataset (left) and splitted by SARS-CoV-2 status (right). **C**. Frequencies of MPC subsets among all MPC across control, mild/moderate (M/M) and severe individuals **D**. Volcano plot showing results of differential gene expression (DGE) analysis performed on all MPC between mild/moderate (right) and severe (left) patients. **E**. Scatter plot between neutrophil and monocyte ISG positive subsets patient by patient. **F**. Correlation matrix using Spearman Rank Correlation between the frequency of all neutrophils and monocytes subtypes in all SARS-CoV-2 negative and SARS-CoV-2 positive patients. (n=32) **G**. Violin plot of ISG signature on all T cells (top) and all B/Plasma cells (bottom) across SARS-CoV-2 status and disease severity. **H**. Scatter plot between neutrophil and CD4 T cell ISG positive subsets patient by patient. **I**. Correlation matrix using Spearman Rank Correlation between the most and the least correlated cell subsets to the Neutrophils ISG positive cells (data include all SARS-CoV-2 negative and positive patients). Statistical significance was assessed using Spearman method (n=32) (G.) Kruskal Wallis test with multiple comparisons (C.). * p-value < 0.05; ** p-value < 0.01; *** p-value < 0.001; **** p-value < 0.0001.

ISG monocyte and ISG neutrophil frequencies were strongly correlated with one another in mild/moderate SARS-CoV-2 positive individuals (**Fig 2E**). Correlating multiple neutrophil subsets versus mononuclear phagocyte subsets across the entire cohort confirmed the strong correlation between ISG neutrophils and ISG monocytes and highlighted a significant correlation between pDCs and NEAT1 neutrophils, which have been previously described in viral infection (30072977) (**Fig 2F**). Similarly, we performed a comprehensive analysis of T cell and B cell frequencies (**Fig S4**) and found again that both cell types are significantly enriched in ISG signatures, specifically in mild/moderate COVID-19 patients (**Fig 2G**). On a patient-by-patient basis, the populations of ISG+ in one compartment correlated with the frequency of ISG expressing cells in another, for example of ISG+ T cells and ISG+ neutrophils, uniquely in mild/moderate patients (**Fig 2H**). Spearman correlation analysis across multiple cell types in all patients thus showed a collection of correlated ISG populations and a second anti-correlated block of other cell populations, notably those expressing S100A12 (**Fig 2I**).

Platelets are the mediators of blood coagulation and their activation can be associated with inflammation and infectious diseases, termed immunothrombosis(*8*). In patients infected with SARS-CoV-2, 25-33% of patients present with thrombocytopenia or thromboembolic events(*9*). We sought to determine how this might be reflected in changes in the expressed genes and concomitantly in the heterotypic cell doublet frequencies in SARS-CoV-2 infections versus other respiratory infections. Our scRNA-seq whole blood data set allowed us to identify and subset platelets based on established platelet signature genes: PPBP, PF4, CLEC1B, RGS10, RGS18 (**Fig 1B/S1C**). Analysis of these platelets revealed six clusters (**Fig 3A/B**), including three subsets (“H3F3B”, “HIST1H2AC”, and “RGS18”) characterized by high expression levels of histone protein-encoding transcripts. Such populations may represent early platelets, based upon the suggestion that these anucleated cells carry transcripts acquired from their parental cells, megakaryocytes, during recent platelet formation(*10*). One by one comparison of these platelet subtypes among healthy controls and patients with mild/moderate and severe disease revealed only minor depletion of HIST1H2AC platelets with disease severity and increase in ACTB platelets—expressing genes associated with cytoskeletal functions—in both mild/moderate and severe patients (**Fig 3C**).

**Figure 3:**
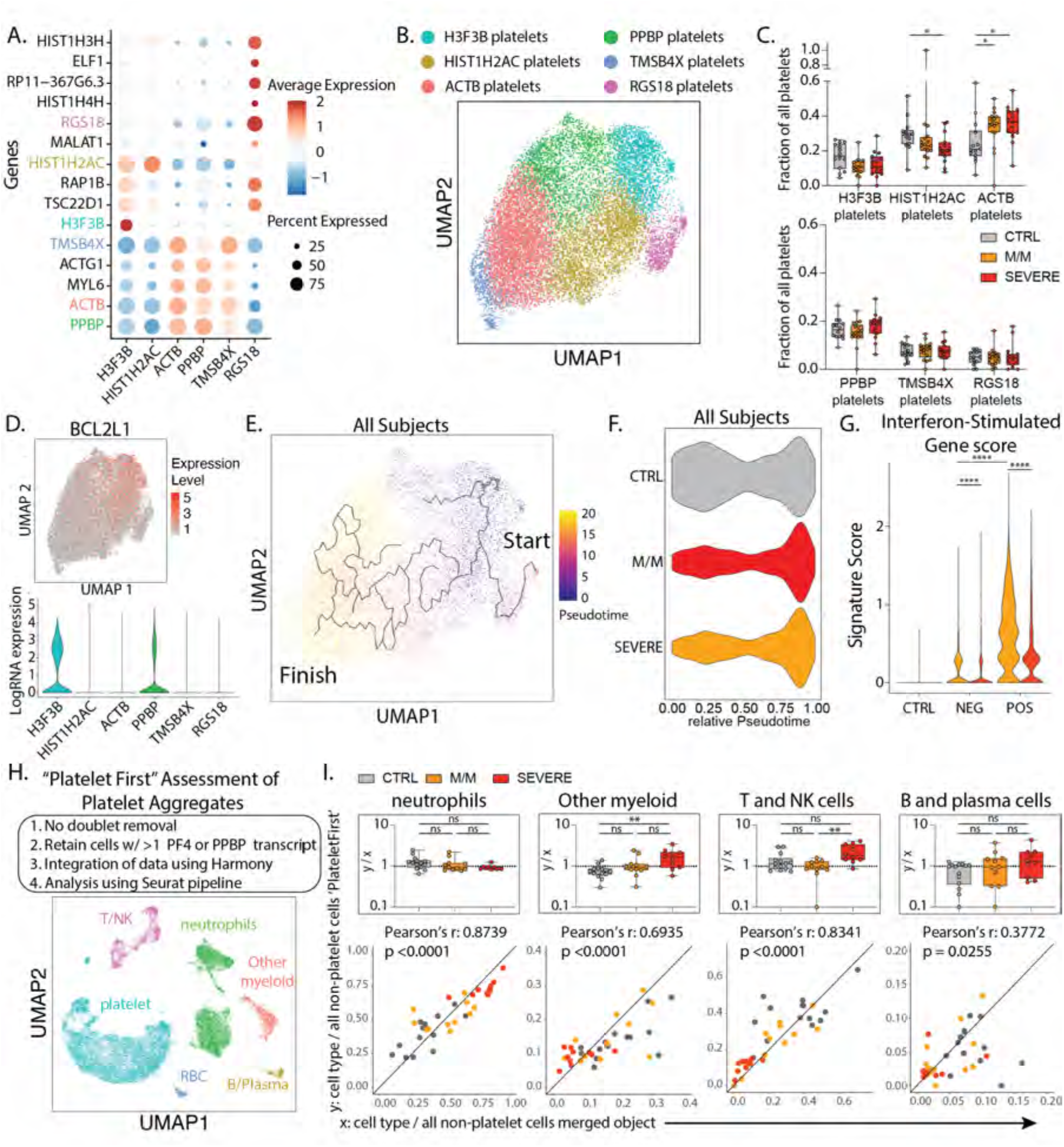
Platelet subtypes and putative platelet aggregates in COVID-19 disease. **A.** Dotplot representation of the top DEG between clusters identified in the platelet subset. **B.** UMAP visualization of 16,903 platelets isolated from the entire dataset showing various subsets, colored distinctly by their identity. **C.** Frequencies of the identified clusters among all platelets in healthy donors and all patients with mild/moderate (M/M) and severe disease. **D.** UMAP visualization of all platelets colored by *BCL2L1* (top) and violin plot of *BCL2L1* expression level across all identified platelet subsets. **E.** UMAP visualization of all platelets with overlay of Pseudotime trajectory. **F.** Violin plot of the relative Pseudotime of all platelets split by healthy donors, mild/moderate and severe patients. **G.** ISG signature score in all platelets across SARS-CoV-2 status and disease severity. **H.** Outline of ‘Platelet First’ assessment to identify platelet aggregates in entire whole blood scRNA-seq data set. UMAP visualization of the 52,757 putative platelet aggregates with specific populations overlaid. **I.** Bottom: Scatter plot of cell type frequency within merged object of entire cohort shown in Figure 1B (*x*-axis) versus same cell type frequency within ‘Platelet First’ object (*y*-axis). The identity line *x=y* is drawn as a reference. Each dot represents a healthy control or SARS-CoV-2 positive patient sample and are color-coded by disease severity. Pearson r correlation coefficient and two-tailed p value are shown for each cell type. Top: Box plots of *y/x-* ratio for each healthy control or patient sample, separated by disease severity. Differences in C. and I. were calculated using a two-way ANOVA test with multiple comparisons. * p.value < 0.05; ** p.value < 0.01; *** p.value <0.001; **** p.value < 0.0001; ns: non-significant.

To compare platelet turnover between disease severity groups, we overlaid the expression of BCL2L1 onto our platelet data set. This transcript encodes the anti-apoptotic protein Bcl-xL, which has been identified as a ‘molecular clock’ for platelet lifetime(*11*). This identified the H3F3B cluster as representing ‘young’ platelets (**Fig 3D**), a result supported by a second signature of transcripts in young, reticulated platelets(*12*) (**Fig S5A**). This population was thus supported as the starting point for a pseudotime analysis, in which the histone-high clusters (H3F3B, HISTH2AC, RGS18) and cytoskeletal protein-high clusters (ACTB, PPBP, TMSB3X) were observed at the start and end of the pseudotime trajectory, respectively (**Fig 3E/S5B**). Plotting the platelet cell frequencies of our cohort along the trajectory suggested that platelets from all patients with disease were broadly overrepresented at the end of the trajectory (**Fig 3F**), supporting a discrete and systemic relative loss of young platelets in all patients as compared to controls. Platelets did not harbor an ISG-specific cell cluster (**Fig S5C**), but akin to myeloid and lymphoid cells, the ISG score in all platelets from mild/moderate patients was increased relative to controls and was comparatively decreased in severe patients, particularly in SARS-CoV-2 infected individuals (**Fig 3G**).

Taking advantage of this unique dataset, we explored the identification of heterotypic aggregates between platelets and non-platelets by using a ‘Platelet First’ approach **(Fig 3H**, see methods). This approach analytically prioritized capture of every cell that was aggregated with a platelet, prior to doublet assessment. The ‘Platelet First’ object revealed the presence of platelet transcripts associated with cells that also bore signatures of other major blood cell types (**Fig 3H/S5D-E**). We separately analyzed a smaller object that included only libraries containing samples pooled prior to cell encapsulation, allowing us to assess inter-patient doublets to conclude that the majority of these aggregates were not formed during the scSeq pipeline (**Fig S5F**). In plotting the blood cell frequencies within the ‘Platelet First’ object against the cell frequencies within the original data set, we found a largely linear relationship between the frequency of a given aggregating population (x-axis) and the frequency with which that cell is found in an aggregate (y-axis) (**Fig 3I**). This first-order linear relationship suggests that, at least in circulating blood, platelets form aggregates indiscriminately with varying other cell types without favoring one or the other. Possibly activated vasculature would provide a cue that would change this pattern of aggregation.

After observing that ISG expression profiles were elevated in every cell type among pateints with mild/moderate disease and COVID-19 but globally reduced with severe COVID-19 disease, we turned our attention to a hypothesis generating holistic approach to data analysis. In an attempt to visualize the global shift in gene expression data across cell types to identify trends with clinical correlates. We first undertook a phenotypic earth mover’s distance (PhEMD) approach (*13*) that identifies and differentiates joint cell frequency patterns to highlight sources of patient-to-patient variation. PhEMD embedding of patients based on their subtype frequencies revealed eight distinct groups of patients (**Fig 4A/S6A**) where progression from A through H represent patients with generally increasing relative frequency of neutrophils, C, D, G and H include patients with relative enrichment in monocytes and E represents patients with an enrichment of ISG neutrophils and mostly consists of SARS-CoV-2 positive patients with mild/moderate disease (**Fig 4B-C**). In contrast, Group G, which is an alternative and ‘severe’ fate for patients is enriched for neutrophil and has a high ratio of cell frequency of S100A12 to ISG neutrophils (**Fig S6A**).

**Figure 4:**
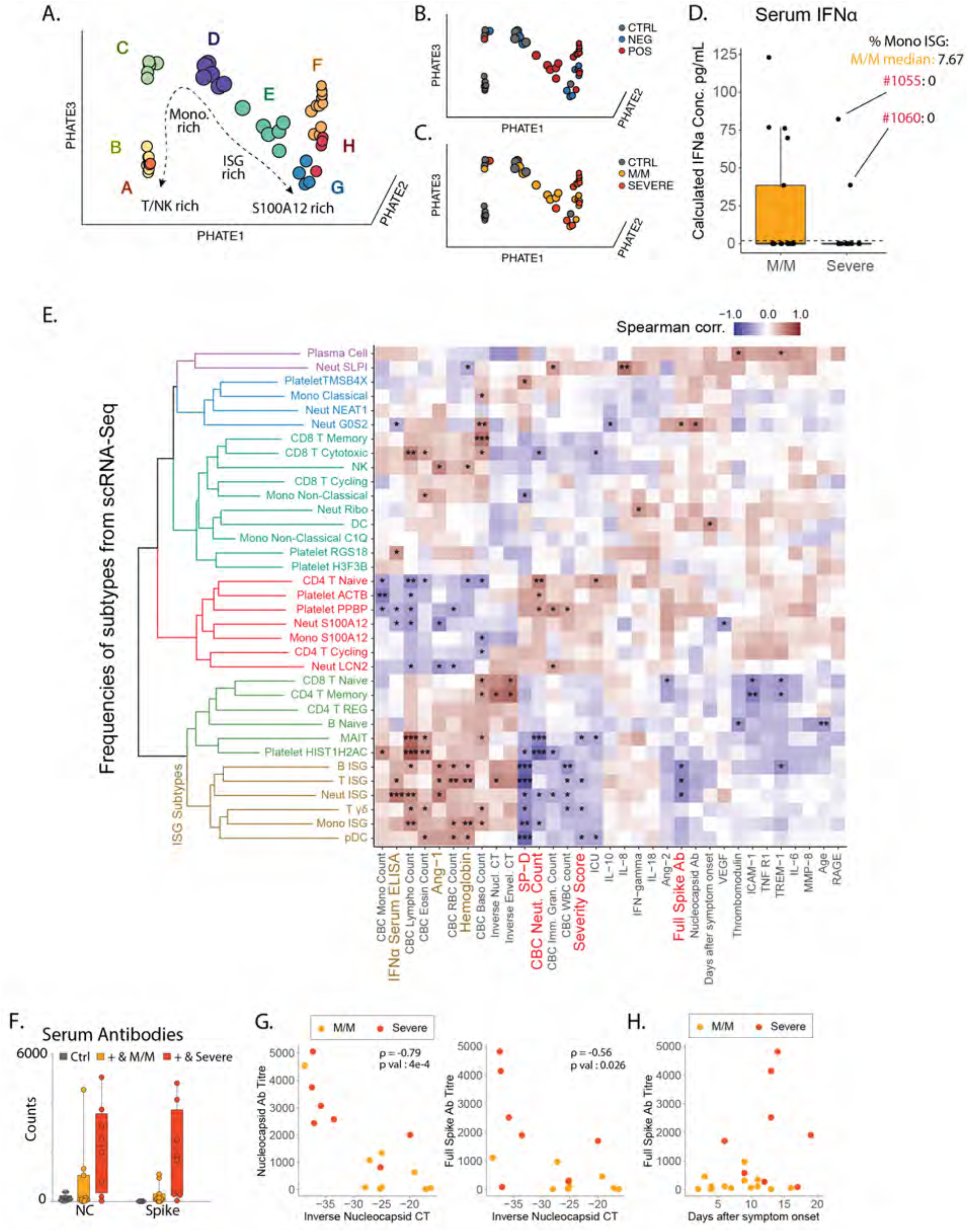
Integrated view of Blood Composition in COVID-19 Patients. **A-B.** 3D PhEMD embedding of all patients, colored by **A**. de novo patient clusters A-H, **B.** SARS-CoV-2 status, and **C**. disease severity. **D.** Measurement of serum IFNα concentration from SARS-CoV-2 positive and negative patients by ELISA. Patients 1055 and 1060 are highlighted in red and their Monocytes ISG frequency from Fig 2C is noted as well as the median for mild COVID-19 mild/moderate patients. **E.** Matrix of Spearman correlation coefficients between all subtype frequencies (out of major cell types, e.g. Neut ISG out of all Neutrophils) obtained from scRNA-Seq versus patient metadata, viral load, Ab titers, and serum analyte levels on a patient-by-patient basis excluding healthy controls. Patients for which data were unavailable were excluded from correlation analysis for each comparison. Variables on both axes were ordered via hierarchical clustering with the computed dendrogram displayed for subtype frequencies. This dendrogram was divided into 6 groupings with the one containing ISG+ subtypes highlighted in brown. Clinical variables generally correlated with severity highlighted in red and anti-correlated in brown. (n for correlation comparisons ranged from n=14 to 32) * p<0.05, ** p<0.005, *** p<0.0005. **F.** Measurement of anti-SARS-CoV-2 antibody levels in serum from patients by Luminex assay (M/M: Mild/Moderate). **G.** Scatter plots showing viral load versus levels of antibody binding SARS-CoV-2 Nucleocapsid and Full Spike protein for patients in the cohort with severity overlaid. Antibody levels are shown as arbitrary units of MFI from Luminex assay while viral load is represented by an inverse CT number from QRT-PCR with target amplification of the SARS-CoV2 Nucleocapsid sequence. Correlation coefficient and significance calculated using Spearman’s method. Patients for which data was unavailable were excluded. (n=16) **H.** Scatterplot for SARS-CoV2 Full Spike protein antibody titers relative to days post symptom onset. Patients for which data was unavailable were excluded. (n=22)

When we examined serum IFNα levels, we found that mild/moderate individuals made more of this cytokine on average as compared to severe patients, which would be consistent with higher levels of ISG cell populations, however there were patient with severe disease individuals who also made high levels of IFNα (**Fig 4D**). To integrate the scRNA-Seq cell populations in the context of other clinical and serum fractions, we constructed a Spearman correlation matrix comparing all subtype frequencies described above with a collection of serum cytokines, antibodies and clinical variables (**Fig 4E**). Following hierarchical clustering of variables, ISG subtypes cluster together and are correlated with serum IFNα concentrations, consistent with these cells arising in response to globally high concentrations of this cytokine as shown previously (3) (**Fig S6B).** ISG-expressing populations are also associated with low severity of COVID-19 illness, lower plasma levels of SP-D (indicative of alveolar epithelial injury) and, only modestly, with IL-6 levels. We also note the absence of significant correlation between ISG populations and days after symptoms onset, indicative of a disease ‘state’ (**Fig 4E** and **Fig S1A**). We further correlated our subtype frequencies against a high-dimensional panel of plasma protein levels (**Fig S6B**) and again observed clustering of most ISG subtypes, which correlated with a host of factors indicative of a strong ISG and Th1 response (CXCL1/6/10/11, TNFB, IL-12B, MCP-2/4), while inversely correlated with others (CCL23, MMP10, HGF). An unexpected anticorrelate of the ISG state was the concentration of serum antibodies against the SARS-CoV-2 Spike and Nucleocapsid proteins.

This anticorrelation was profound (**Fig 4F**) and we considered it a paradox that severe patients have higher levels of potentially neutralizing antibodies. This is in apparent contradiction with a previous report showing that viral load is associated with severity and mortality in COVID-19(*14, 15*) a difference which could be explained by the fact that these studies focus on patients with high mortality, which was a very rare event in our cohort **(Sup Table S1)**. Both antibody specificities were anticorrelated with the viral load as assessed from nasal swabs (**Fig 4G**) consistent with though not definitive for being neutralizing. As increased antibody titers and decreased viral load have been reported to be a feature of later disease stage(*16*), we considered the hypothesis that mild/moderate disease – characterized by high frequency of ISG+ signatures – would simply precede severe disease states. However, antibody titers in severe patients are consistently higher compared to mild/moderate patients even two weeks beyond symptom onset **(Fig 4H)**, and only one of our 11 mild/moderate patients would go on to exhibit a severe disease **(Fig S1A)**. Moreover, we observed no statistical correlation between days of onset and the presence of ISG+ cell populations **(Fig 4E).** These elements would seem to argue against a simple time relationship between mild/moderate and severe states.

Returning to the profound antibody reactivity and the global loss of ISG populations even in the presence of serum IFNα **(Fig 4D)**, we hypothesized that phenotypic differences in our two groups of COVID-19 patients might also be mirrored or influenced or by systemic factors such as those carried in the blood and affecting all cell populations. We thus first asked whether serum from severe patients contained antibodies against ISG-expressing cells by directly applying serum to healthy peripheral blood mononuclear cells (PBMCs) from heathy individuals that were first induced to express IFITM3 (part of our ISG signature, encoding a protein that blocks viral entry (*6*)) by culturing them with IFNα for 24 hours, followed by flow cytometry analysis of monocytes and lymphocytes **(Fig S7A)**. While we observed only low levels of serum IgG binding in serum from 1/9 mild/moderate patients, sera from 5/7 patients with severe COVID-19 illness displayed significant binding (**Fig 5A/S7B**). Staining was more pronounced on monocytes as compared to T and B cells which were largely stained with similar frequencies when using mild/moderate or severe serum (**Fig S7C**). When examining specificities in those patients that did not stain ISG-differentiated cells directly, we found that serum from patient 1050 produced antibodies that were specific for IFNα (**right inset Fig5A**), consistent with a very recent study (*17*) that also found these in approximately 12% of COVID patients. This finding nevertheless does not explain patients lacking ISG cells despite presence of IFNα in their serum (**Fig 4D**).

**Figure 5:**
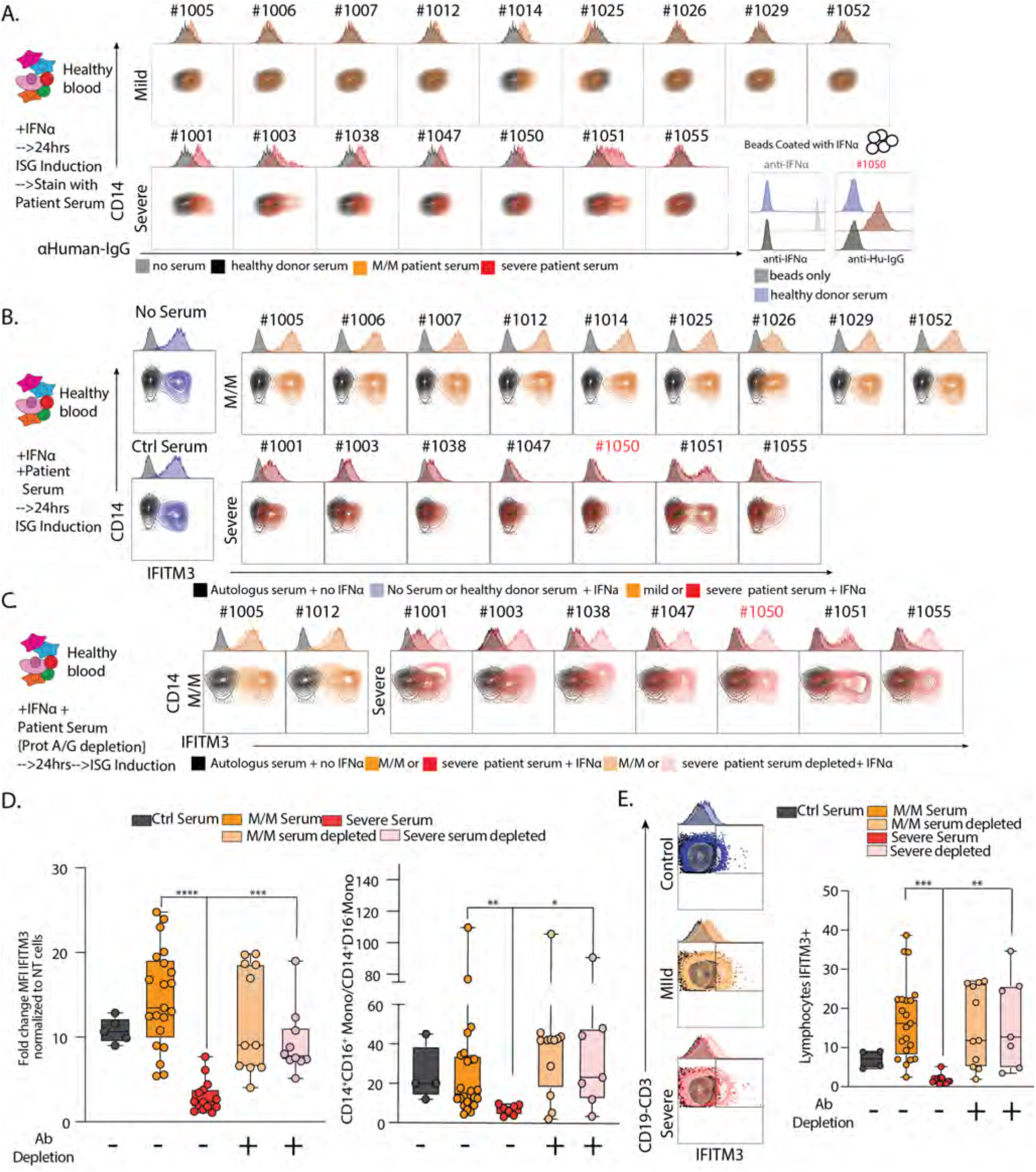
Neutralization of ISG induction by Antibodies from Severe COVID-19 Patients. **A.** Contour plots and histograms of CD14 Monocytes from healthy blood cultured with IFNα to induce expression of ISGs and stained with serum from heathy donor, mild/moderate (M/M) or severe SARS-CoV-2 positive patients with secondary staining with α-human IgG. Bottom right: histogram of beads coated with IFNα and stained with an antibody raised against IFNα or serum from severe SARS-CoV-2 positive patient #1050 or healthy donor. Black histograms represent non coated beads. **B-C.** Contour plots and histograms of CD14 Monocytes from healthy blood cultured with IFNα and serum from heathy donor, mild/moderate or severe SARS-CoV-2 positive patient quantifying levels of intracellular IFITM3 staining. **C.** Mild/Moderate (light yellow) or Severe (pink) sera were pre-treated with protein G/A before incubation with PBMC. **D.** Box plot of IFITM3 induction in CD14 monocytes (left) and intermediate to classical monocytes ratio (right) from 2 different experiment and 2 different healthy donors. **E.** Left: Contour plots and histograms of pooled CD3+/CD19+ lymphocytes from healthy blood cultured with IFNα and serum from heathy donor, mild/moderate or severe SARS-CoV-2 positive patients. Mild/moderate (light yellow) or Severe (pink) sera were pre-treated with protein G/A before incubation with PBMC to deplete antibodies. Right: Box plot of IFITM3 induction in lymphocytes. Differences in D. and E. were calculated using a two-way ANOVA test with multiple comparisons. * p.value < 0.05; ** p.value < 0.01; *** p.value <0.001; **** p.value < 0.0001; ns: non-significant.

We then asked whether factors in the serum of severe patients affect the induction of the ISG signature gene pattern, including IFITM3, in response to culture with IFNα. We thus added patient serum at 10% into the IFNα stimulation conditions and found that, whereas control serum or serum from mild/moderate patients had no effect on differentiation as measured by either IFITM3 level or the frequencies of CD14+CD16^+^ intermediate monocytes produced (**Fig 5B/5D and Fig S7D**), all severe patient serum tested had profound effects on differentiation, varying from absolute block to partial inhibition.

To test whether these effects on IFN-induced production of ISG cell populations were in fact mediated by antibodies, we pre-adsorbed patients’ sera with Protein A/G beads to deplete them. This antibody depletion restored both IFITM3 induction and the total yield of interferon-stimulated monocyte cells (**Fig 5C/D S7D**). A similar block of ISG signature generation in response to IFNα was observed for other populations including lymphocytes, showing that this effect was global and similarly mediated by antibodies in serum severe patient (**Fig 5E/S7D-E**). Moreover, we were able to confirm inhibition of ISG cell population generation by serum from severe patients, in a second validation cohort (data not shown).

Taken together, this shows that an antibody response in severe patients targets ISG cell populations and their generation. In our cohort, this general effect of antibody manifests in all severe patients, whereas antibodies against the cytokine IFNα itself were seen only in one of seven patients, a similar frequency recently reported(*17*). Antibodies in many of these patients have direct specificity for determinants on the surface of ISG monocytes. The molecular specificities of these other antibodies are likely to be many and varied based on the differing patterns in this relatively small sample set and it will remain to be determined how and why tolerance is broken to the ISG pathway, in the course of infection. One likely candidate for modulation of B cell response is direct infection of monocytes by SARS-CoV-2. *In vitro* incubation of the virus with healthy cells indeed results in intracellular expression of both IFITM3 (indicating activation of this program) and spike protein **(Fig S7F).** If, in an early immune response, the ISG program is preferentially presented alongside the proteins from the pathogen, and the immune system of the patient is not already self-tolerant to those ISG proteins due perhaps to a lifelong absence of their profound expression, tolerance to those cells and those proteins may be broken. Conversely, as inflammatory monocytes normally restrict antibody generation (*18*), their infection and lysis by virus may in turn release overexuberant B cell responses to many antigens, not just those that are newly produced during the infection. Regardless, this work suggests that targeting overexuberant and autoreactive B cells with drugs such as rituximab (*19*) or through introduction of IVIG(*20*), perhaps alongside introduction of convalescent serum-derived antibodies to provide ongoing viral protection, may be an avenue to defeat the global suppression of protective ISG mediated viral immunity.

## Acknowledgements

We thank all members of the Krummel Lab and ImmunoX for discussion and guidance while developing this study. We would like to thank Dr Gaia Andreoletti for discussion and guidance on computational analysis. This work was supported by funds from the UCSF ImmunoX Initiative and funds from the NIH (R01 AI52116-S1 (MFK) 3U19AI077439-13S1 (DJE), NHLBI R35 HL140026(CC)). K.H.H. is supported by the American Cancer Society Postdoctoral Fellowship (#133078-PF-19-222-01-LIB). A.R. is a Cancer Research Institute Irvington Postdoctoral Fellow supported by the Cancer Research Institute (Award # CRI2940). This project has been made possible in part by grant number 2019-202665 from the Chan Zuckerberg Foundation. The data reported in this manuscript are in the main paper and in the supplementary materials. The authors declare no competing financial interests.

## Author Contributions

AJC, TC, NFK, KHH, AR, AAR, WSC, SC, MFK performed and provided supervision of experiments, generated and analyzed data, and contributed to the manuscript by providing figures and tables. AJC, TC, NFK, KHH, AR, AAR, MFK had full access to all of the data in the study and take responsibility for the integrity of the data and the accuracy of the analyses. AJC, TC, NFK, KHH, AR, AAR, WSC, performed computational analyses of the data. AJC, TC, KHH, VC, NWC, DK, GCR, AS, JT, KJHG, PMS, WSZ, DSL, YS, were part of the early morning COVID-19 processing team who performed the whole blood single cell sequencing. The UCSF COMET Consortium included all other scientist involved in the generation of data included in this study. CL provided viral load obtained from nasal swab of COVID-19 patient. CJY, ML and SC provided blood from healthy donors and helped with PBMC preparation. RY, MM,LR, AL, CRZ, RPL performed assays of SARS infection and serum antibodies and cytokines. MW and MM oversaw studies of serum cytokines and antibodies from the COVID-19 patient cohort and provided intellectual input. PGW, DJE, GKF, CC, CL, CH, MFK are leadership members of the COMET study and were actively involved in the establishment of the pipeline and patient consent and the direction of projects. PGW, CC, CH, AW and SC actively participated in patient and control cohort enrollment. AJC, TC, NFK, KHH, AR, AAR, WSC, MFK wrote the manuscript. AJC, AAR and MFK supervised the project. All authors edited and critically revised the manuscript for important intellectual content and gave final approval for the version to be published.

## Supplementary Materials

### Supplemental Table 1

**Supplementary Table 1:** COVID-19 Whole Blood Study Cohort. Patients were enrolled as described in Material and Methods and blood was collected from 4:30AM rounds on the first day after admission. Healthy controls were from volunteers who fasted overnight and were collected between 6am and 9am. Data on individuals is shown along with average and standard deviations.

| COMET ID | SARS-CoV-2 Status | Disease Severity | Age | Gender (1=MALE, 2 =F) | Days between symptoms and sampling (* = asymptomatic) | Other Infection(s) | ICU during hospital stay | Days under mechanical Ventilation | Discharged |
| --- | --- | --- | --- | --- | --- | --- | --- | --- | --- |
| 1008 | NEG | Mild/Moderate | 35.7 | 1 | 20 | Bacterial | No | 0 | Yes |
| 1016 | NEG | Mild/Moderate | 63.5 | 2 | 4 | None | No | 0 | Yes |
| 1017 | NEG | Mild/Moderate | 65.2 | 2 | 9 | None | No | 0 | Yes |
| 1045 | NEG | Mild/Moderate | 82.6 | 1 | NA | None | Yes | 0 | Yes |
| 1049 | NEG | Mild/Moderate | 68.8 | 2 | 1 | Bacterial | No | 0 | Yes |
| 1062 | NEG | Mild/Moderate | 75.8 | 2 | NA | Bacterial | No | 0 | Yes |
| **COVID Negative Mild n=6** | **Average Data:** |  | **65** | **1.7** | **9** |  |  |  |  |
|  | **SD:** |  | **16** |  | **8** |  |  |  |  |
| 1009 | NEG | Severe | 67 | 1 | NA | Bacterial | Yes | 5 | yes |
| 1010 | NEG | Severe | 50.7 | 2 | 9 | Bacterial | Yes | 5 | Yes |
| 1020 | NEG | Severe | 78.6 | 1 | 5 | Bacterial | Yes | 2 | Yes |
| 1021 | NEG | Severe | 77 | 2 | 2 | Bacterial | Yes | 5 | Yes |
| 1046 | NEG | Severe | 49.6 | 1 | 3 | Bacterial | Yes | 1 | Yes |
| **COVID Negative Severe n=5** | **Average Data:** |  | **65** | **1.4** | **5** |  |  |  |  |
|  | **SD:** |  | **14** |  | **3** |  |  |  |  |
| 1005 | POS | Mild/Moderate | 83.4 | 1 | 9 | None | No | 0 | Yes |
| 1006 | POS | Mild/Moderate | 71.9 | 2 | 16 | None | No | 0 | Yes |
| 1007 | POS | Mild/Moderate | 55.3 | 1 | 13 | Bacterial | No | 0 | Yes |
| 1012 | POS | Mild/Moderate | 41.1 | 1 | 3 | None | No | 0 | Yes |
| 1014 | POS | Mild/Moderate | 58.6 | 1 | 4 | None | Yes | 0 | Yes |
| 1025 | POS | Mild/Moderate | 73.2 | 1 | NA* | None | No | 0 | Yes |
| 1026 | POS | Mild/Moderate | 45.5 | 2 | 6 | None | No | 0 | Yes |
| 1029 | POS | Mild/Moderate | 85 | 2 | 2 | Viral | No | 0 | Yes |
| 1044 | POS | Mild/Moderate | 73.9 | 1 | NA* | None | No | 0 | Yes |
| 1052 | POS | Mild/Moderate | 37.9 | 2 | 4 | Viral | No | 0 | Yes |
| 1054 | POS | Mild/Moderate | 25.8 | 1 | NA* | None | No | 0 | Yes |
| **COVID Positive Mild n=11** | **Average Data:** |  | **59** | **1.4** | **7** |  |  |  |  |
|  | **SD:** |  | **20** |  | **5** |  |  |  |  |
| 1001 | POS | Severe | 34.8 | 2 | 12 | Viral | Yes | 10 | Yes |
| 1002 | POS | Severe | 79.6 | 1 | 4 | Bacterial | Yes | 7 | Yes |
| 1003 | POS | Severe | 43.6 | 2 | 13 | Viral | Yes | 5 | Yes |
| 1031 | POS | Severe | 44.2 | 1 | 20 | Viral | Yes | 28 | Yes |
| 1038 | POS | Severe | 61.4 | 2 | 14 | None | Yes | ND | Yes |
| 1047 | POS | Severe | 48.3 | 2 | 6 | Viral + Bacterial | Yes | 28 | Yes |
| 1050 | POS | Severe | 55 | 1 | 13 | Bacterial | Yes | 29 | Yes |
| 1051 | POS | Severe | 63.3 | 1 | 19 | Bacterial | Yes | 33 | Yes |
| 1055 | POS | Severe | 62.4 | 1 | 4 | Viral | Yes | 31 | Yes |
| 1060 | POS | Severe | 44.1 | 1 | 6 | Bacterial | Yes | 5 | Yes |
| **COVID Positive Severe n=10** | **Average Data:** |  | **54** | **1.4** | **11** |  |  |  |  |
|  | **SD:** |  | **13** |  | **6** |  |  |  |  |
| ICC_0001 | CTRL | CTRL | 39 | 1 | NA | NA | NA | NA | NA |
| ICC_0003 | CTRL | CTRL | 24 | 1 | NA | NA | NA | NA | NA |
| ICC_0666 | CTRL | CTRL | 45 | 2 | NA | NA | NA | NA | NA |
| ICC_1084 | CTRL | CTRL | 34 | 1 | NA | NA | NA | NA | NA |
| ICC_1117 | CTRL | CTRL | 41 | 1 | NA | NA | NA | NA | NA |
| ICC_1367 | CTRL | CTRL | 52 | 1 | NA | NA | NA | NA | NA |
| ICC_3231 | CTRL | CTRL | 28 | 2 | NA | NA | NA | NA | NA |
| ICC_3954 | CTRL | CTRL | 37 | 2 | NA | NA | NA | NA | NA |
| ICC_4096 | CTRL | CTRL | 31 | 1 | NA | NA | NA | NA | NA |
| ICC_4117 | CTRL | CTRL | 59 | 2 | NA | NA | NA | NA | NA |
| ICC_4319 | CTRL | CTRL | 30 | 1 | NA | NA | NA | NA | NA |
| ICC_4444 | CTRL | CTRL | 34 | 2 | NA | NA | NA | NA | NA |
| ICC_4799 | CTRL | CTRL | 40 | 2 | NA | NA | NA | NA | NA |
| ICC_4923 | CTRL | CTRL | 34 | 1 | NA | NA | NA | NA | NA |
| **Healthy Controls n=14** | **Average Data:** |  | **38** | **1.4** | **NA** |  |  |  |  |
|  | **SD:** |  | **9** |  | **NA** |  |  |  |  |

### Supplementary Figures S1–S5

**Supplementary Figure 1:**
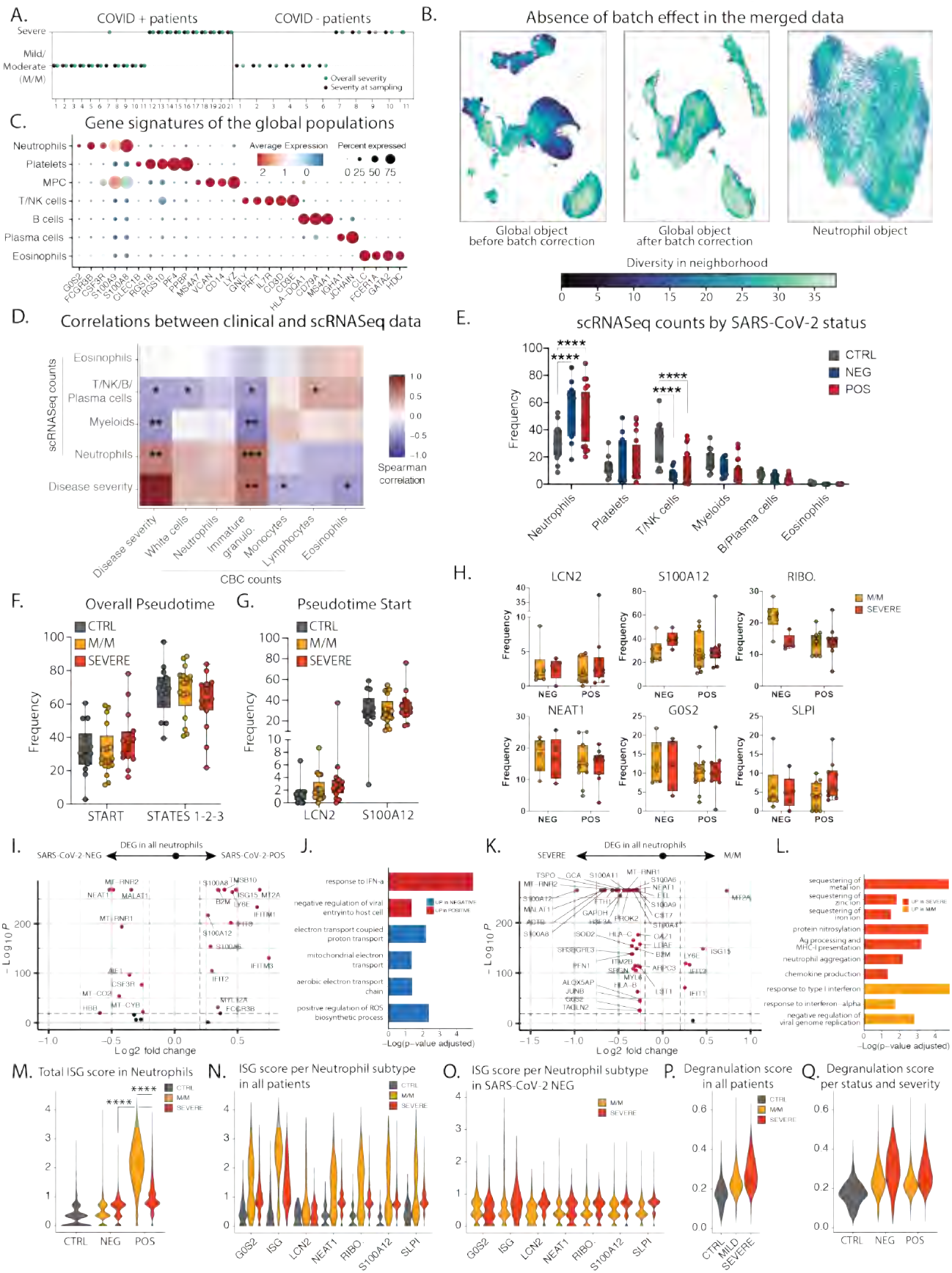
**A**. Patient symptoms plot: symptom at day of sampling (first day of admission to the hospital) is represented in black, while symptom based on the entire course of hospitalization is in green. In the rest of the manuscript we categorized patient into mild or severe cases based on all the entire course of hospitalization (green). **B**. Quantification of the batch effect using neighbor diversity score in the global object UMAP before (left) and after (middle) batch correction, along with the neutrophil (right) UMAP plot, as in Fig1B and Fig1D, using the diversity in neighborhood method. **C**. Dotplot representation of landmark genes expressed by global populations in Fig1B. **D**. Spearman’s correlation comparison between between disease severity and population frequencies calculated from 10X scRNAseq analyses (10X) or complete blood cell counts (CBC). Patients for which CBC counts were unavailable were excluded. Significance was calculated using Spearman’s method. * p value<0.05; ** p value <0.05; *** p value<0.005 (n=29) **E**. Frequency of the global populations in Fig1B among all cells across SARS-CoV-2 status. **F and G**. Frequencies of the neutrophil subsets among all neutrophils across control, mild/moderate (M/M) and severe individuals at the overall start/late states of the trajectories (F) or at specific early stages of the pseudotime (G). **H**. Frequency of the LCN2, S100A12, RIBO., NEAT1, G0S2 and SLPI neutrophils among all neutrophils across SARS-CoV-2 status and disease severity. **I to L**. Volcano plots showing DEG (I and K) and bar plots showing GO term enrichment from these DEG (J and L) between all neutrophils from either SARS-CoV-2 positive vs negative patients (I and J) or mild/moderate vs severe patients (K and L) **M to Q.** Scores of ISG signature (M to O) and neutrophil degranulation (P and Q) in either all neutrophils across control, mild/moderate an severe patients (P), all neutrophils across SARS-CoV-2 status and disease severity (M and Q) or specific neutrophil subtypes across severity in either all patients (N) or only SARS-CoV-2 negative patients (O). Statistical significance was assessed using a two-way ANOVA test with multiple comparisons for panels E, F and H, and using a Wilcoxon test for panel M. * p.value < 0.05; ** p.value < 0.01; *** p.value < 0.001; **** p.value < 0.0001.

**Supplementary Figure 2:**
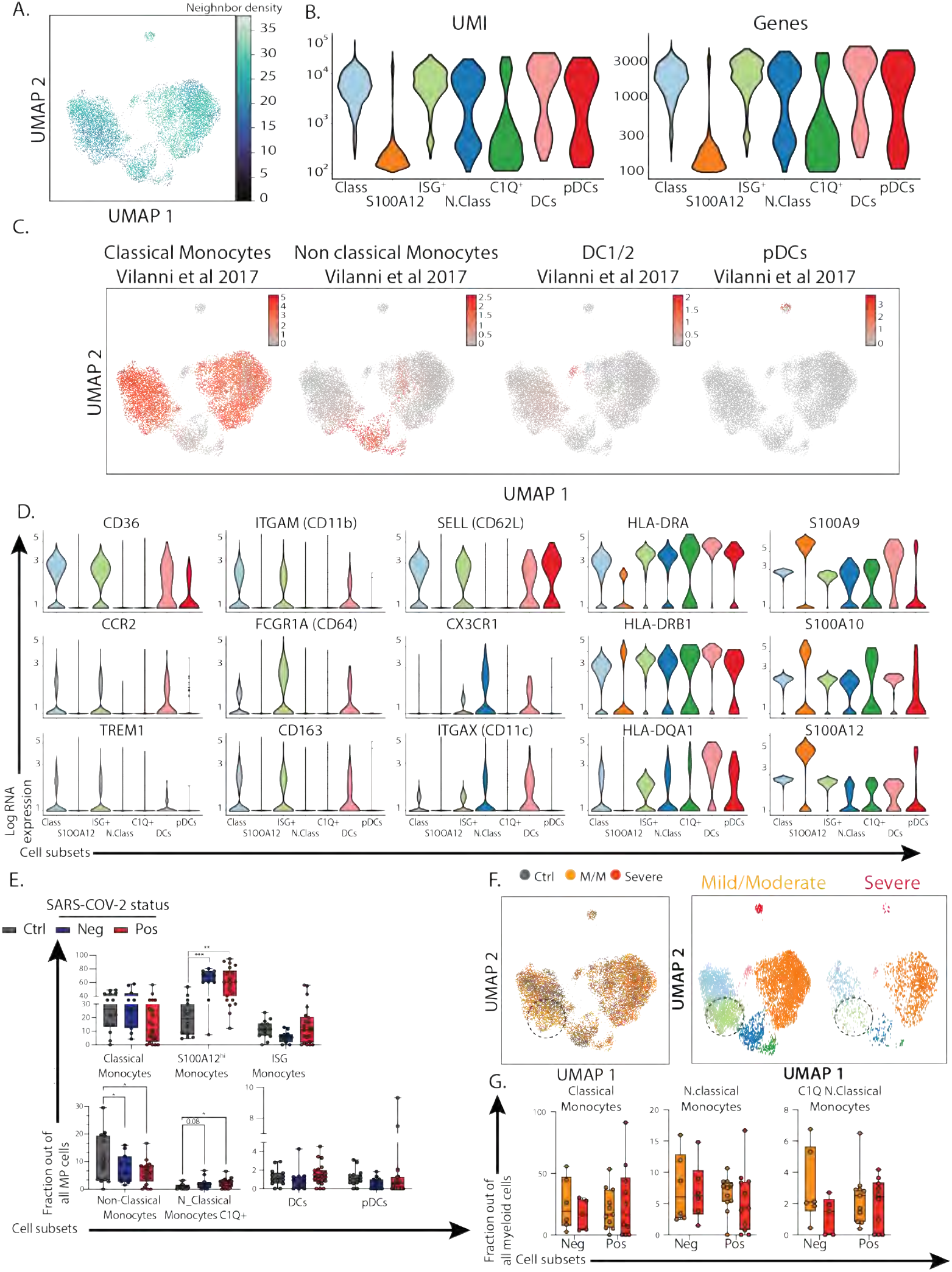
**A**. Quantification of the batch effect before and after batch correction using neighbor diversity score in the mononuclear phagocytic cells (MPC) object from UMAP plot in Fig2C, using the diversity in neighborhood method. **B**. Violin plot of number of unique genes (bottom) and number of unique molecules (top) detected from Single cell sequencing for each cluster identified in the MPC dataset. **C**. Overlay of previously described blood mononuclear phagocytic cell signature from healthy individual on MPC from UMAP plot in Fig2C. **D**. Violin plots of canonical genes previously described as expressed by blood MPC for each for each cluster identified in the MPC dataset. **E**. Frequencies of the MPC subsets among all MPC across control, SARS-CoV-2 negative and SARS-CoV-2 positive individuals. **F**. UMAP visualization of the 19,289 MPC colored (left) and split by (right) by disease severity. **G**. Frequencies of the classical monocytes, cycling monocytes, non-classical monocytes and C1Q+ non classical monocytes among all MPC across SARS-CoV-2 negative and SARS-CoV-2 positive individuals split it by disease severity. Statistical significance was assessed using a two-way ANOVA test with multiple comparisons. * p.value < 0.05; ** p.value < 0.01; *** p.value < 0.001; **** p.value < 0.0001.

**Supplementary Figure 3:**
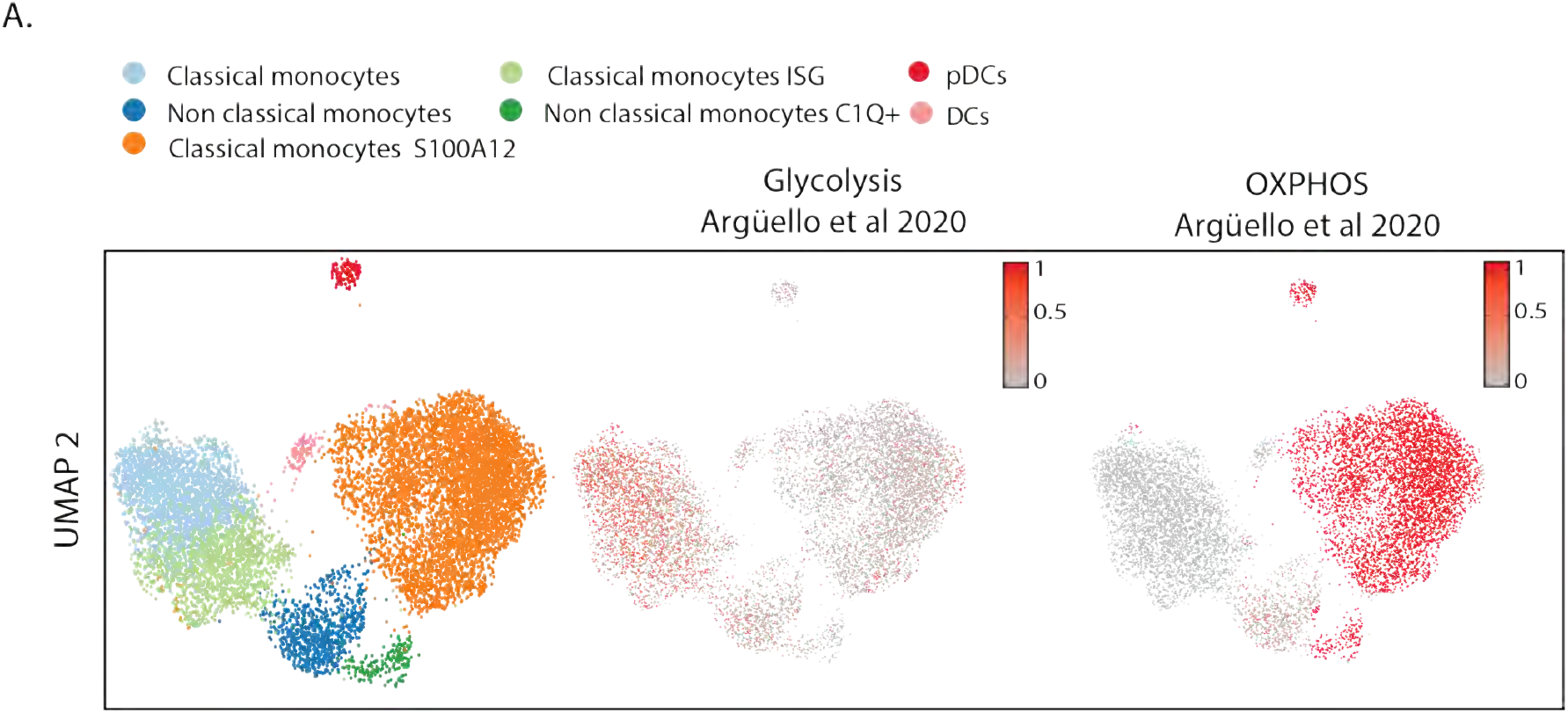
**A.** Overlay of previously described glycolytic and oxidative phosphorylation gene signature on mononuclear phagocytic cells (MPC) from UMAP plot in Fig2C.

**Supplementary Figure 4:**
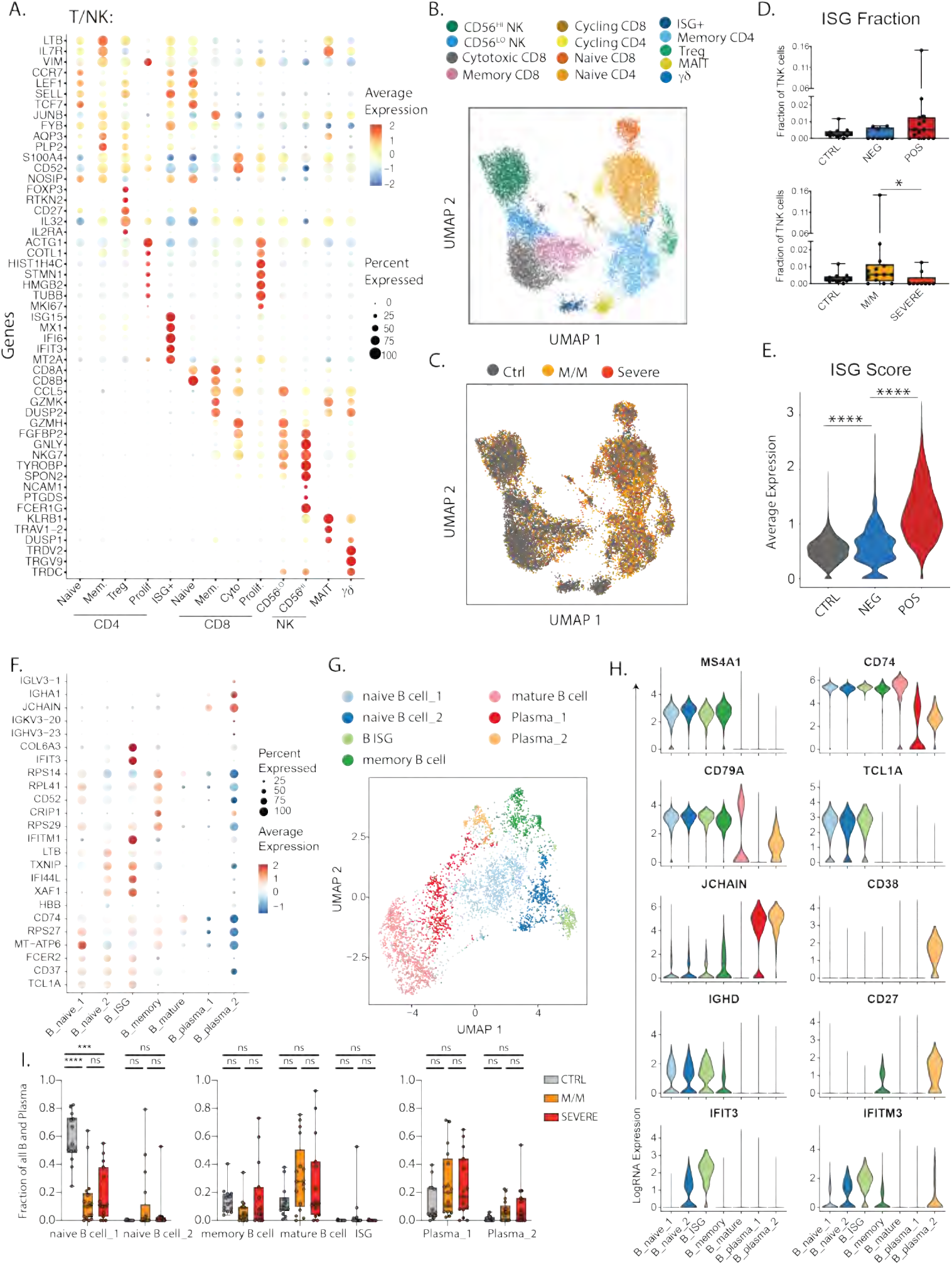
**A.** Dotplot representation of the top DEG between clusters identified in the T and NK cell subset. **B.** UMAP visualization of 16,708 T and NK cells in the entire dataset showing various subsets, colored distinctly by their identity. **C.** Overlay of the above UMAP of all T and NK cells, colored by disease severity underlining the lack of batch effects while merging the datasets from all patients. **D**. Abundance of the Interferon-stimulated-gene (ISG) + subset among all T and NK cells in healthy donors, SARS-CoV-2 negative and SARS-CoV-2 positive patients (top) and in healthy donors and patients with mild/moderate (M/M) and severe disease (bottom). **E.** ISG signature score between healthy controls, SARS-CoV-2 negative and SARS-CoV-2 positive patients. **F.** Dotplot representation of the top DEG between clusters identified in the B and plasma cell subset. **G.** UMAP visualization of 4,380 B and plasma cells isolated from the entire dataset showing various subsets, colored distinctly by their identity. **H.** Violin plots of canonical genes previously described as expressed by B and plasma cells for each identified cluster. **I.** Frequencies of the identified clusters among all B and plasma cells in healthy donors and patients with mild/moderate and severe disease. Differences in D. and E. were calculated using Kruskal-Wallis non-parametric ANOVA with multiple comparisons. * p <0.05 and **** p< 0.001. Differences in I. were calculated using a two-way ANOVA test with multiple comparisons. * p.value < 0.05 and **** p.value < 0.0001. ns, non-significant.

**Supplementary Figure 5:**
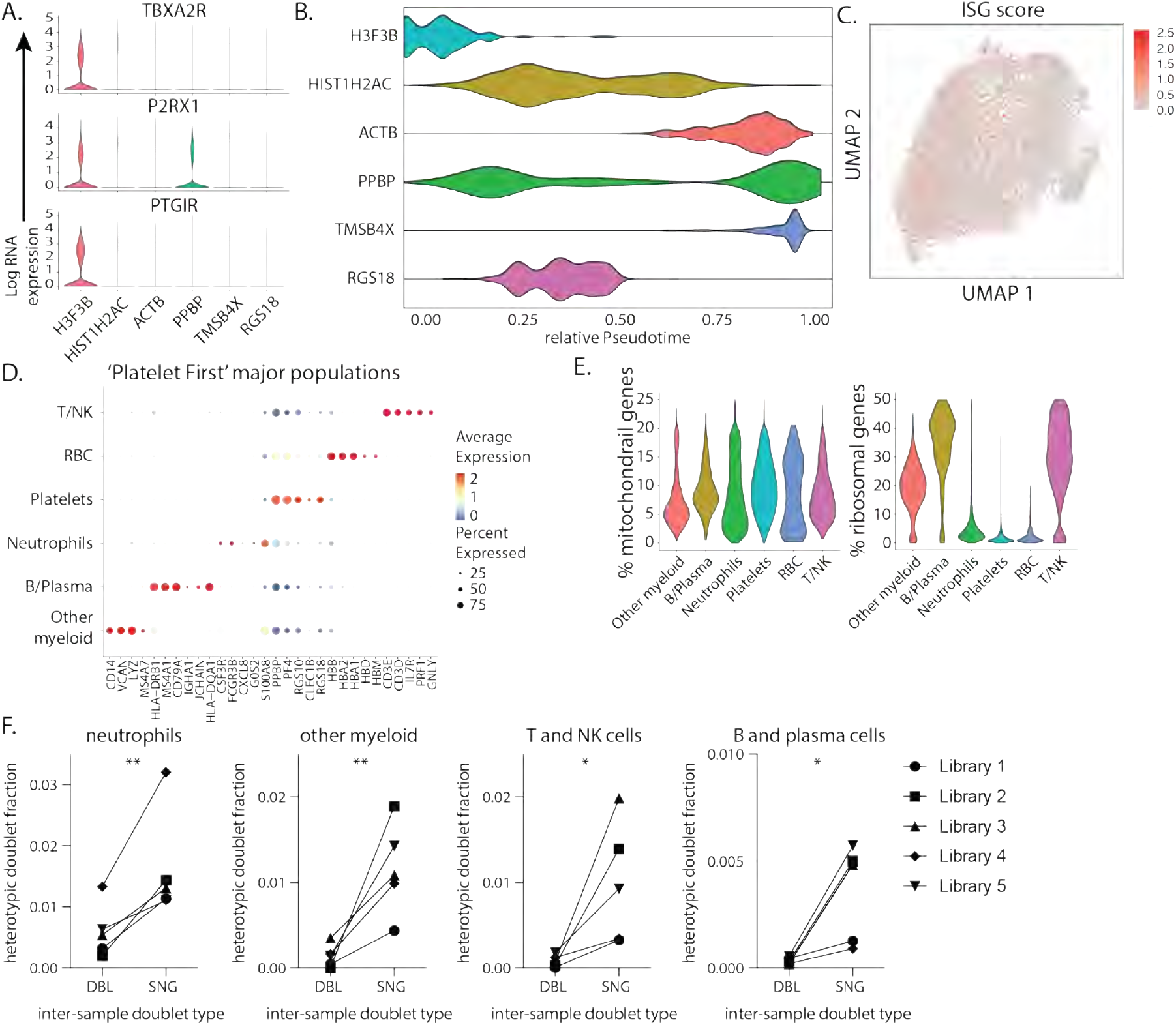
**A.** Violin plots of genes identifying young, reticulated platelets (*1*) in the platelet dataset. **B.** Violin plots of the relative pseudotime of each platelet cell subset present in Figure 3B. **C.** UMAP visualization of all platelets colored by ISG score. **D.** Dotplot representation of the top DEG between clusters identified in the ‘Platelet First’ object. In this object (see also Fig 3) no doublet removal filtering step was applied to include all heterotypic cellcell aggregates (Step 1). This was followed by retaining all cells with >1 platelet-specific transcripts PF4 or PPBP (Step 2). Step 2 guaranteed analysis of cell events and aggregates containing platelets. Identically to our original data set in Figure 1B, integration of data was done using Harmony (Step 3), and the ‘Platelet First’ object was then analyzed using the Seurat v3 pipeline (Step 4). **E.** Violin plots of the percentage of mitochondrial and ribosomal genes within clusters identified in the ‘Platelet First’ object. **F.** Inter-sample doublet rates in inferred platelet-involved heterotypic doublets show that platelet aggregates occur in vivo. DBL, doublet. SNG, singlet. Differences in F. were calculated using a onesided Student’s t test * p.value < 0.05 and ** p.value < 0.01.

**Supplementary Figure 6:**
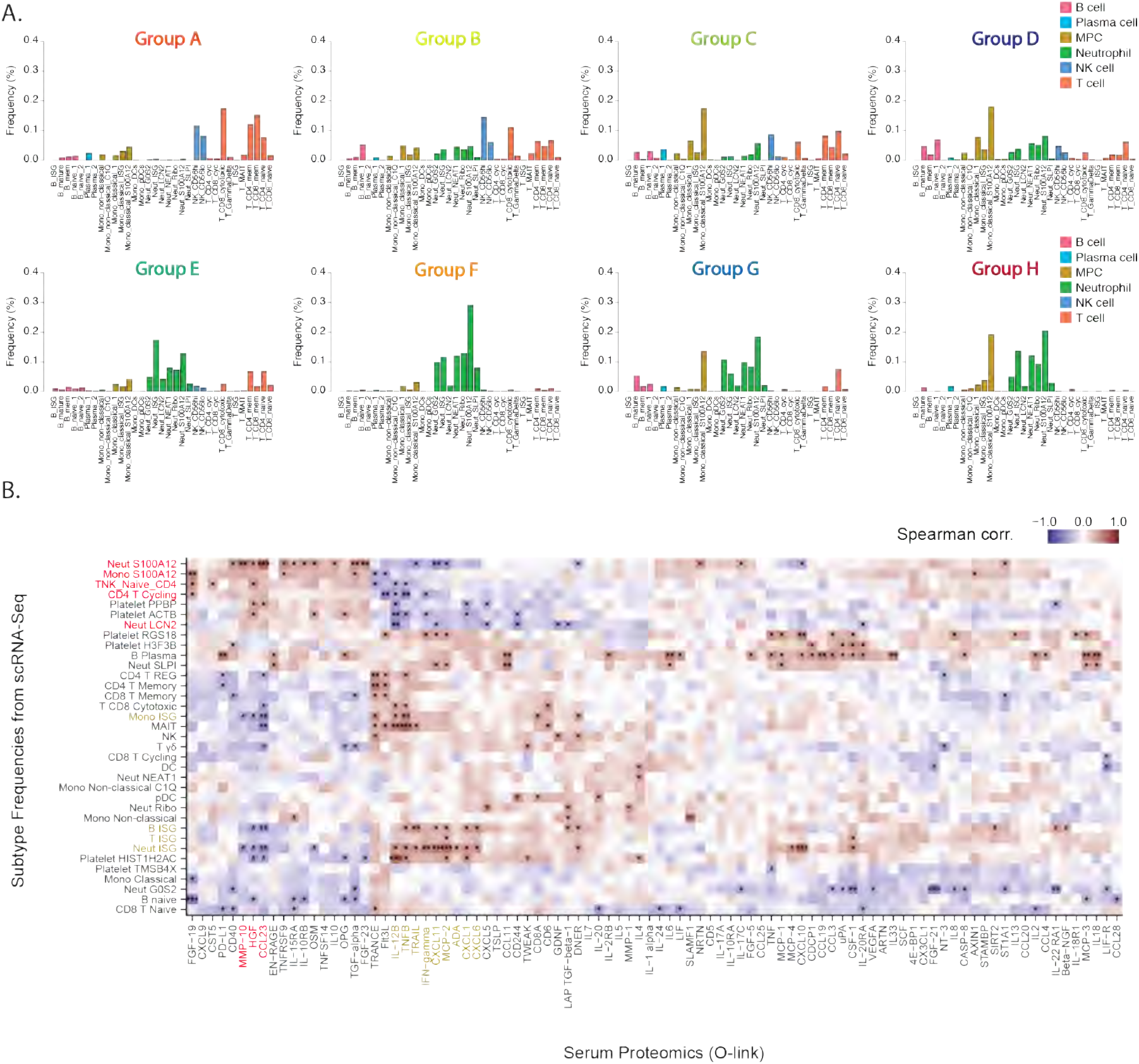
Identifying Immune System Correlates in SARS-CoV-2 positive and negative Patients Data. **A.** Cell fraction histograms representing bin-wise mean of relative frequency (i.e., cell fraction) of each immune cell subtype for all patients in a given group, colored as described in Fig4A. **B.** Matrix of Spearman correlation coefficients between all subtype frequencies (out of major celltypes e.g. Neut ISG represents % out of all Neutrophils) obtained from scRNA-Seq versus protein analyte abundance in plasma as measured using Olink assay on a patient-by-patient basis excluding healthy controls. Patients for which data were unavailable were excluded from correlation analysis for each comparison. Variables on both axes were ordered via hierarchical clustering. ISG subtypes and protein levels strongly correlated with their frequency highlighted in brown. Subtypes and proteins strongly anti-correlated with ISG+ subtypes highlighted in red. (n=31 for all comparisons) * p<0.05, ** p<0.005, *** p<0.0005.

**Supplementary Figure 7.**
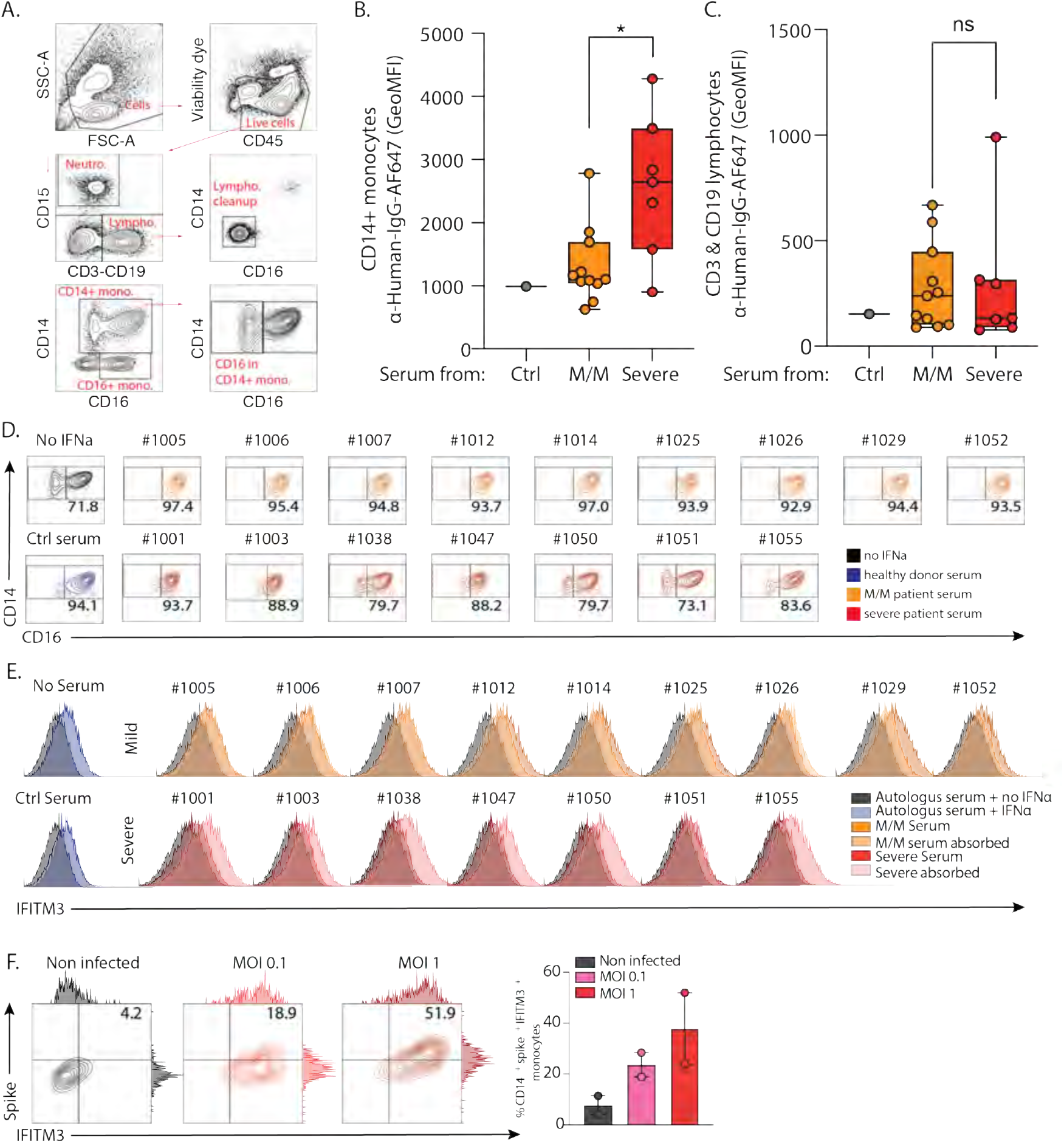
Quantification of Serum Staining and ISG Generation in the Presence of Patient Serum. **A.** Gating strategy for PBMCs to identify different subpopulations. **B.** Geometric MFI of serum staining on CD14+ monocytes treated with IFNα, quantifying data in Figure 5A. **C.** Geometric MFI of serum staining of CD3+ and CD19+ lymphocytes, following treatment with IFNα, quantifying data in Figure 5A. **D.** Modulation of intermediate to classical CD14 monocytes transition by mild/moderate (orange) and severe (red) patient serum. Each plot represents a single serum sample. Representative experiment from three independent trials and two different healthy PBMC donors. **E.** Histograms CD3+ CD19+ lymphocytes from healthy donor cultured with IFNα and serum from heathy donor (blue), mild/moderate (orange) and severe (red) SARS-CoV-2 positive patients. Mild/Moderate (light yellow) or Severe (pink) sera were pre-treated with protein G/A before incubation with PBMC. Each plot represents a single serum sample. Representative experiment from two independent trials and two different healthy PBMC donors. **F.** Contour plots and histograms of CD14+ monocytes from healthy donor co-cultured with SARS-COV-2 virus for 48h at 0.1 (pink) and 1 (red) MOI. Differences in B. and C. were calculated using a two-way ANOVA test with multiple comparisons. * p.value < 0.05; ** p.value < 0.01; *** p.value <0.001; **** p.value < 0.0001. ns, non-significant.

## Material and Methods

### Patients, participants, severity score, and clinical data collection

Patients admitted to the Hospital of the University of California with known or presumptive COVID-19 were screened within 3 days of hospitalization. Patients, or a designated surrogate, provided informed consent to participate in the study. This study includes a subset of patient enrolled between April 8 and May 1 in the COMET (COVID-19 Multi-immunophenotyping projects for Effective Therapies; https://www.comet-study.org/) study at UCSF. COMET is a prospective study that aims to describe the relationship between specific immunologic assessments and the clinical courses of COVID-19 in hospitalized patients. Healthy donors (Ctrl) were adults with no prior diagnosis of or recent symptoms consistent with COVID-19. This analysis includes samples from participants who provided informed consent directly, via a surrogate, or otherwise in accordance with protocols approved by the regional ethical research boards and the Declaration of Helsinki. For inpatients, clinical data were abstracted from the electronic medical record into standardized case report forms. We used both a severity score at the time of sampling and at the end of hospitalization (Fig S1A). In both cases, severity assessment was based on three main parameters: level of care, need for mechanical ventilation, and time under mechanical ventilation. Mild/moderate patients are floor/ICU patients who did not require mechanical ventilation during their time of hospitalization and spent no more than 1 day in ICU. Severe patients are patients who required intensive care and mechanical ventilation (typically 5 days or more). Therefore, our cohort is composed of 21 COVID-19 positive patients (11 mild/moderate and 10 severe), 11 COVID-19 negative patient (6 mild/moderate and 5 severe), and 14 Healthy participants. Information on age, sex, type of infection, days of on onset, viral load, and CBC count are listed in Table S1. The study is approved by the Institutional Review board: IRB# 20-30497.

### Isolation of blood cells and processing for scRNA-seq

ScRNA-seq was performed on fresh whole blood in order to preserve granulocytes. Briefly, peripheral blood was collected into EDTA tubes (BD, catalog no. 366643). Whole blood was prepared by treatment of 500μL of peripheral blood with RBC lysis buffer (Roche, 11-814-389-001) according to manufacturer’s procedures. Cells were then counted and 15.000 cells per individual were directly loaded in the Chromium™ Controller for partitioning single cells into nanoliter-scale Gel Bead-In-Emulsions (GEMs) following manufacturer’s procedures (10x genomics). Some samples were pooled together (at 15,000 cells/ sample) prior to GEM partitioning. Single Cell 5’ reagent kit v5.1 was used for reverse transcription, cDNA amplification and library construction of the gene expression libraries (10x Genomics) following the detailed protocol provided by 10x Genomics. Libraries were sequenced on an Illumina NovaSeq6000 using 28 cycles for R1 and 98 cycles for R2. All samples were encapsulated, and cDNA was generated within 6 hours after blood draw.

### Bulk RNASeq library preparation for Genotyping

RNA was extracted from aliquots of 250K Peripheral Blood Mononuclear Cells (PBMCs) utilizing the ZYMO Research Quick RNA MagBead kit (R2133) on a Thermofisher KingFisher Flex system following manufacturer’s procedures. RNA integrity was inspected with Agilent Fragment Analyzer. Ribosomal and hemoglobin depleted total RNA-sequencing library were created using FastSelect (Qiagen cat#: 335377) and Tecan Universal Plus mRNA-Seq (0520-A01) with adaptations for automation of a Beckmen BioMek FXp system. Libraries were subsequently normalized and pooled for Illumina sequencing using a Labcyte Echo 525 system available at the Center for Advanced Technology at UCSF. The pooled libraries were sequenced on an Illumina NovaSeq S4 flow cell lane with paired end 150bp reads.

### Computational Processing for Genotyping

Sequencing reads were aligned to the human reference genome and Ensembl annotation (GRCh38 genome build, Ensembl annotation version 95) using STAR v2.7.5c (PMID: 23104886) with the following parameters: --outFilterType BySJout --outFilterMismatchNoverLmax 0.04 --outFilterMismatchNmax 999 --alignSJDBoverhangMin 1 --outFilterMultimapNmax 1 --alignIntronMin 20 --alignIntronMax 1000000 --alignMatesGapMax 1000000. Duplicate reads were removed and read groups assigned by individual for variant calling using Picard Tools v2.23.3 (http://broadinstitute.github.io/picard/). Nucleotide variants were identified from the resulting bam files using the Genome Analysis Tool Kit (GATK, v4.0.11.0) following the best practices for RNA-seq variant calling (PMID: 25431634; PMID: 21478889). This include splitting spliced reads, calling variants with HaplotypeCaller (added parameters: --dont-use-soft-clipped-bases -standcall-conf 20.0), and filtering variants with VariantFiltration (added parameters: -window 35 - cluster 3 --filter-name FS -filter FS > 30.0 --filter-name QD -filter QD < 2.0). Variants were further filtered to include a list of high quality SNP for identification of the subject of origin of individual cells by removing all novel variants, maintaining only biallelic variants with MAF greater than 5%, a mix missing of one individual with a missing variant call at a specific site and requiring a minimum depth of two (parameters: --max-missing 1.0 --min-alleles 2 --max-alleles 2 --remove-indels --snps snp.list.txt --min-meanDP 2 --maf 0.05 --recode --recode-INFO-all –out).

### Data pre-processing of 10x Genomics Chromium scRNA-seq data

Sequencer-obtained bcl files were demultiplexed into individual sample the mkfastqs command on the Cellranger 3.0.2 suite of tools (https://support.10xgenomics.com). Feature-barcode matrices were obtained for each sample by aligning the raw fastqs to GRCh38 reference genome (annotated with Ensembl v85) using the Cellranger count. Raw feature-barcode matrices were loaded into Seurat 3.1.5(*2*) and genes with fewer than 3 UMIs were dropped from the analyses. Matrices were further filtered to remove events with greater than 20% percent mitochondrial content, events with greater than 50% ribosomal content, or events with fewer than 100 total genes. The cell cycle state of each cell was assessed using a published set of genes associated with various stages of human mitosis (*3*).

### Inter-sample doublet detection

Libraries containing samples pooled prior to loading were processed using Freemuxlet (https://github.com/statgen/popscle), the genotype-free version of Demuxlet(*4*), to identify clusters of cells belonging to the same patient via SNP concordance. Briefly, the aligned reads from Cellranger were filtered to retain reads overlapping a high-quality list of SNPs obtained from the 1000 Genomes Consortium (1KG)(*5*). Freemuxlet was run on this filtered bam using the 1KG vcf file as a reference, the input number of samples/pool as a guideline for clustering groups of cells by SNP concordance, and all other default parameters. Cells are classified as singlets arising from a single library, doublets arising from two or more libraries, or as ambiguous cells that cannot be accurately assigned to any existing cluster (due to a lack of sufficient genetic information). Clusters of cells belonging to a unique sample were mapped to patients using their individual Freemuxlet-generated genotype, and ground truth genotypes per patient identified via bulk RNASeq. The pairwise discordance between inferred and ground-truth genotypes was assessed using the bcftools gtcheck command(*6*). Ambiguous, and doublet events were filtered from the major analysis (see platelet-first analysis).

### Data quality control and Normalization

The filtered count matrices were normalized, and variance stabilized using negative binomial regression via the scTransform method offered by Seurat(*7*). The effects of mitochondrial content, ribosomal content, and cell cycle state were regressed out of the normalized data to prevent any confounding signal. The normalized matrices were reduced to a lower dimension using Principal Component Analyses (PCA) and the first 30 principal coordinates per sample were subjected to a non-linear dimensionality reduction using Uniform Manifold Approximation and Projection (UMAP). Clusters of cells sharing similar transcriptomic signal were identified using the Louvain algorithm, and clustering resolutions varied between 0. 6 and 1.2 based on the number and variety of cells obtained in the datasets. Clusters were loosely grouped into major cell types (T/NK, B/Plasma, mononuclear phagocytes, Neutrophil, Platelet, and Erythrocytes) using a curated list of 5 genes per cell type (5-gene signature) and the Seurat AddModuleScore function. Briefly, genes in the library are binned into one of 12 bins based on average expression in the dataset. The average expression of the genes in each signature are compared to a background list of randomly selected from the bins and used to generate a score per cell for each signature.

### Intra-sample heterotypic doublet detection

All libraries were further processed to identify heterotypic doublets arising from the 10X sample loading. Processed, annotated Seurat objects were processed using the DoubletFinder package (*8*). Briefly, the cells from the object are modified to generate artificial duplicates, and true doublets in the dataset are identified based on similarity to the artificial doublets in the modified gene space. The prior doublet rate per library was approximated using the information provided in the 10x knowledgebase (https://kb.10xgenomics.com/hc/en-us/articles/360001378811) and this was corrected to account for homotypic doublets using the per-cluster numbers in each dataset. Heterotypic doublets were removed from the major analysis (see platelet-first analysis).

### Data integration and Batch correction

The individual processed objects per library were filtered to remove Erythrocyte contamination. The raw and log-normalized counts per library were then pruned to retain only genes shared by all libraries. Pruned counts matrices were merged into a single Seurat object and the batch (or library) of origin was stored in the metadata of the object. The log-normalized counts were reduced to a lower dimension using PCA and the individual libraries were aligned in the shared PCA space in a batch-aware manner (Each individual library was considered a batch) using the Harmony algorithm (*9*). The resulting Harmony components were used to generate a batch corrected UMAP, and to identify clusters of transcriptionally similar cells. Clusters were broadly labeled based on the 5-gene signature (Fig S1) using a modified, bootstrapped version of the Seurat AddModuleScore to account for the numerous sequencing batches in our dataset. The modified function ran the AddModuleScore on random subsets of the data (subsampling rate = 0.6) 10 times and averaged the score to provide a stable score per signature. A Seurat object was generated for each broad cell type containing clusters scoring highly for that cell type. Each broad cell-type object was subjected to the same harmony analysis to generate batch-aware log normalized counts that were used for visualization and subtype identification. To visualize the effect of harmonizing our single cell data, we identified the library diversity in the neighborhood of every cell on the plot. The neighborhood of a cell is defined as the collection of n nearest neighbors in the UMAP space (where n = sqrt(total cells)), pruned to retain cells lying within the 90th percentile of all calculated neighbor distances. The diversity is the set of all libraries represented within the neighborhood.

### Differential expression tests and cluster marker genes, cluster annotation and volcano plot

Differential gene expression (DGE) tests were performed on log-normalized gene counts using the Poisson test (with a latent batch variable to account for multiple library preparations) provided by the FindMarkers/FindAllMarkers functions in Seurat. Genes with > 0.35 log-fold changes, an adjusted p value of 0.05 (based on Bonferroni correction), at least 25% expressed in tested groups, were regarded as significantly differentially expressed genes (DEGs). Cluster marker genes were identified by applying the DE tests for upregulated genes between cells in one cluster to all other clusters in the dataset. Top ranked genes (by log-fold changes) from each cluster of interest were extracted for further illustration. The exact number and definition of samples used in the analysis are specified in the legend of Figure 1, 2 and 3 and summarized in Table S1. The neutrophils, mononuclear phagocytic cells, T cells, B cells and platelets subtypes were identified by comparing cluster marker genes with public sources referenced in the text. The R package EnrichR were used to generate volcano plot from differential gene expression using FindMarkers function in Seurat.

### Platelet First scSeq Analysis

To identify the differential coagulation of platelets, we reintroduced heterotypic doublets to each library and filtered them to extract cells expressing at least 1 UMI of PF4 or PPBP, both platelet-specific marker genes. The raw and log-normalized counts per library were integrated using Harmony and processed as above. Broad cell types were identified using the score generated with the bootstrapped AddModuleScore and the per-sample rate of platelet aggregation with each cell types was inferred to be the fractions of cell counts in this dataset to the fractions of cell counts in the overall analysis. Significance testing was conducted using a non-parametric Kruskal-Wallis test with multiple comparisons.

### Monocle analysis

Raw counts from the Individual cell-specific were used to create a monocle3 (*10–12*) cell_data_set object, and the batch-corrected PCA and UMAP embeddings were imported directly from the Seurat object. Each cell-specific trajectory was inferred by reverse embedding the UMAP coordinates using the DDRTree method. The root cell states for the trajectory in monocytes and neutrophils were chosen based on literature, and for platelet cell based on the signature list defined in Figure S1. Relative pseudotime was obtained through a linear transformation relative to the cells with the lowest and highest pseudotimes ((*1-min_pseudotime)/max_pseudotime*).

### Generation of gene expression scores

ISG and Degranulation scores were generated by taking the mean of log-normalized expression for a particular set of genes related to the specific pathway or phenotype. The following gene lists were used to generate the scores-ISG: *MT2A, ISG15, LY6E, IFIT1, IFIT2, IFIT3, IFITM1, IFITM3, IFI44L, IFI6, MX1, IFI27;* Degranulation: 486 genes from Neutrophils degranulation GO term #GO:0043312. To visualize the distribution of these scores across cells, we binarized the distribution of the score at the 75th percentile and overlaid on the calculated UMAP coordinates.

### Correlation plots and heatmap visualization

Correlation coefficients used in variable against variable comparisons were calculated using Spearman’s method to avoid assumption of linearity. Significance testing of correlation was performed with the following tests: Spearman’s for continuous v. continuous, Kruskal-Wallis or Wilcoxon rank sum test for categorical v. continuous depending on number of categories and Fisher’s exact for categorical v. categorical comparisons. Unless otherwise specified, variables on both axes were hierarchically clustered based on the distance matrix computed from the correlation coefficient.

### Embedding a low-dimensional representation of patients using PhEMD

PhEMD was employed to generate a three-dimensional embedding of patients based on their immune cell profiles (*13*). Briefly, PhEMD first generates a reference map of cell subtypes, then uses Earth Mover’s Distance (EMD) to compute pairwise dissimilarities between patients (incorporating patient-to-patient differences in cell fractions of each cell subtype as well as intrinsic dissimilarities between subtypes based on the cell subtype reference map), and finally applies a dimensionality reduction technique to the patient-to-patient distance matrix to generate a final embedding of patients. The Seurat implementation of 3D Uniform Manifold Approximation and Projection (UMAP) was used to map the cell-subtype space using the Harmony batch-corrected components as input and a “min.dist” parameter of 0.4, and cell subtypes (i.e., clusters) were defined as described in the “Data quality control and Normalization” section of Methods (*2, 9*). Dissimilarity between each pair of cell subtypes was defined as the distance between the centroids (in UMAP space) of all cells assigned to the two respective subtypes. PHATE was applied to the EMD patient-to-patient distance matrix to generate the final 3D embedding of patients (*14*).

### Luminex Assay for Antibody Titer

Highly immunogenic linear regions of the SARS-CoV-2 proteome were isolated by ReScan and conjugated to Luminex beads as previously described (*15*). Briefly, high concentration T7 phage stocks displaying immunodominant epitopes of the S, N and ORF3a proteins were propagated and grown to high (>10^11^ PFU/mL) titer then were each conjugated to unique bead IDs according to manufacturer’s Antibody Coupling Kit instructions (Luminex). Whole N protein (RayBiotech) beads were conjugated similarly using manufacturer instructions with 5μg of protein per 1 million beads. For other whole protein Luminex-based beads, MagPlex-Avidin Microspheres (Luminex) were coated with either the S protein RBD (residues 328-533) or the trimeric S protein ectodomain (residues 1-1213). All beads were blocked overnight before use and pooled on day of use. 2000-2500 beads per ID were pooled per incubation with patient serum at a final dilution of 1:500 for 1 hour, washed, then stained with an anti-IgG pre-conjugated to phycoerythrin (Thermo Scientific, #12-4998-82) for 30 minutes at 1:2000. Primary incubations were done in PBST supplemented with 2% nonfat milk and secondary incubations were done in PBST. Beads were processed in 96 well format and analyzed on a Luminex LX 200 cytometer. Median Fluorescence Intensity from each set of beads within each bead ID were retrieved directly from the LX200 and log transformed after normalizing to the mean signal across two intra-assay negative controls (glial fibrillary acidic protein (GFAP) and Tubulin phage peptide conjugated beads).

### Luminex Assay for Serum Cytokines

Soluble proteins were quantified in EDTA anticoagulated plasma using the Luminex^®^ multiplex platform (Luminex, Austin TX) with custom-developed reagents (R&D Systems, Minneapolis, MN), as described in detail ((*16*) or single-plex ELISA (R&D Systems, Minneapolis, MN). Analytes quantified using the Luminex^®^ multiplex platform were read on the MAGPIX^®^ instrument and raw data were analyzed using the xPONENT^®^ software. Analytes quantified using single-plex ELISA were read using optical density. Values outside the lower limit of quantification were assigned a value of 1/3 of the lower limit of the standard curve for analytes quantified by Luminex and 1/2 of the lower limit of the standard curve for analytes quantified by ELISA.

### O-link Assay for Serum Factors

Circulating proteins were measured in plasma using the multiplexed Proximity Extension Assay (PEA) from Olink Proteomics AB (Uppsala, Sweden). 20 μL each of plasma collected from the COMET patient cohort (21 COVID-19 positive, 13 COVID-19 negative, and 14 healthy individuals) were analyzed using the Olink^®^ Target 96 Inflammation panel, which is a set of 92 inflammation-related protein biomarkers. Plasma for all samples regardless of COVID-19 status were inactivated using a final concentration of 1% (v/v) Triton-X-100 solution over 2 hours. Data from the analyzed protein biomarkers is presented as Normalized Protein eXpression (NPX) values, an arbitrary unit on a log2 scale.

### ELISA Method for Serum IFNα Measure

IFN-α levels were quantified from serum by an ELISA (catalog numbers 41115 for IFN-α; PBL Assay Science). ELISA was performed according to the manufacturer’s instructions with minor modifications. Briefly, an 8-point standard curve was prepared and diluted in sample buffer. Serum was also diluted by adding 80-90 ul serum to 30 ul sample buffer (depending on availability of serum). Samples were prepared in duplicates, whereas standards were prepared in triplicates. Initial incubation was performed for 20 hours at 4C. Antibody was added and incubated at 4C overnight. HRP was added and incubated 1hr at room temperature, TMB substrate was added for 30min and incubated in the dark at room temp, stop solution was added and samples were read using a SpectraMax M2 Microplate Reader (Molecular Devices) at 450nm. For analysis, a 4-parameter logistic fit was applied to OD values of the standards after background subtraction. Samples with ODs below blank samples were considered as 0 pg/ml IFN-α.

### PBMC co-culture experiment with patient serum and flow cytometry analysis

PBMCs were isolated from EDTA-anticoagulated whole blood from healthy donors using Polymorphprep (Alere Technologies), and resuspended in culture medium (RPMI 1640 + 10% FBS). For detection of neutralization of interferon stimulation, autologous serum or clinical study participant sera (10 μl) were plated with IFNα (Stemcell IFN alpha-2A; final concentration of 1 pg/ml) in a total volume of 100μl before addition of 4×105 PBMCs. After incubation for 24 hours, PBMCs were assayed for IFNα-induced IFITM3 upregulation and CD14/CD16 levels and fractions by flow cytometry. After surface staining and addition of fixable live/dead violet dye (ThermoFisher; #L34955), intracellular detection of IFITM3 was done using the eBioscience Foxp3 / Transcription Factor Staining Buffer Set (ThermoFisher; #00-5523-00) and following the manufacturer’s instructions. For autoantibody assays, PBMCs were cultured with media or 1-100 pg/ml IFNα for 38-46 hours. Samples were harvested and unconjugated AffiniPure Fab Fragment Goat anti-human IgG (H+L) (Jackson Immunoresearch; #109-007-003) and Human TruStain FcX block (BioLegend; #422302) were used to block pre-bound antibodies and Fc receptors. After washing with fluorescence-activated cell sorting (FACS) buffer (2% fetal bovine serum, 1 mM EDTA, PBS), PBMCs were then stained for surface markers 30 min on ice. After staining incubation, cells were washed 3x times with FACS buffer (1500 rpm, 5 min, 4°C) and incubated with 5μl autologous or clinical study participant sera for 90 min on ice. After washing the cells with FACS buffer, cell-bound antibodies were detected using an AffiniPure Donkey anti-human IgG-Alexa Fluor 647 antibody (Jackson Immunoresearch; #709-605-149), which was incubated with the cells for 30 min on ice. Cells were washed again and resuspended in 1 μg/ml DAPI solution for live/dead discrimination. The following antibodies were used for flow cytometric analysis: anti-human CD3-BB700 (clone SK7; BD Biosciences; #566575), anti-human CD14-BV711 (clone MSE2; BioLegend; #301838), anti-human CD15-BV786 (clone W6D3; BD Biosciences; #741013), anti-human CD16-BV605 (clone 3G8; BioLegend; #302040), anti-human CD19-BB700 (clone SJ25C1; BD Biosciences; 566396), anti-human CD45-APCeFluor780 (clone HI30; ThermoFisher; 47-0459-42), anti-human IFITM3-AlexaFluor 647 (clone EPR5242; Abcam; ab198573).

### Bead ELISA

10^7^ 5um Sulfate latex polystyrene beads (Thermo Fisher) were resuspended in 1ml of PBS to which 1ug of proteins (BSA, huIFNα (Stemcell IFN alpha-2A) were added to bind by passive absorption over 1 hr on ice. Beads were washed 1x in PBS and blocked with 1ml of blocking buffer (PBS containing 1mM EDTA and 2%FCS) for one hour. Beads were spun and resuspended in 1ml of blocking buffer and 10ul (10^5^ beads) were moved to individual tubes to which 5ul of sera was added followed by incubation for 1hr on ice). These were washed, resuspended in 50ul of blocking buffer containing Goat anti-human IgG-Alexa Fluor 647 (Jackson Immunoresearch) and incubated for 1hr on ice followed by a final wash and analysis by flow cytometry.

### SARS-CoV-2 detection by PCR

PCR testing for SARS-CoV-2 was carried out on respiratory specimens mixed 1:1 in DNA/RNA Shield (Zymo Inc) using an in-house Clinical Laboratory Improvement Amendments (CLIA) validated assay at the UCSF Clinical Microbiology Laboratory. PCR primers targeted the SARS-CoV-2 envelope (E) and RNA-dependent RNA polymerase (RdRp) genes plus human RNAse P as a positive control.

### SARS-CoV-2 infection of PMBC

SARS-CoV-2 isolate USA-WA1/2020 was provided by Dr. Melanie Ott and propagated in Vero E6 (ATCC CRL-1586) cells in Dulbecco’s Modified Eagle Medium (UCSF Cell Culture Facility) supplemented with 10% FBS. Vero E6 cells were infected with the SARS-CoV-2 virus for 72h at 37C and 5% CO2. The supernatant was collected and viral titer was quantified using a plaque assay in Vero E6 cells. All work was done under BSL3 conditions. PBMC were infected with SARS-CoV-2 virus at a multiplicity of infection (MOI) 0.1 or 1 for 72 hours. Cells were then harvested and stained with fixable Live/Dead Zombie NIR (Biolegend) in PBS followed by fixation with 4% paraformaldehyde for 1 hour. Intracellular staining to Spike (SARS-CoV-2 Spike S1 Antibody, Rabbit mAb (SinoBiological) and IFITM3 was subsequently performed using eBioscience Foxp3/Transcription Factor Staining Buffer Set (Thermo Fisher Scientific) followed by surface antigen staining.

### Statistical Analysis and Data visualization

Statistical analyses were performed using GraphPad prism or the R software package. Null hypotheses between two groups were tested using the non-parametric Mann-Whitney test to account for non-normal distribution of the data. Likewise, for multiple groups, comparisons were made by two-way ANOVA or non-parametric Kruskal–Wallis test followed by multiple comparisons. The specific statistical tests and their resultant significance levels are also noted in each figure legend. The R packages Seurat, ggplot2 (version 3.1.0) (Wickham, 2016) GraphPad Prism and Adobe Illustrator were used to generate figures.

### Data Resources and Code Sharing

Raw Gene expression matrices will be available on GEO at the time of publication. Scripts used to process all data will be shared on Github along with relevant clinical information for each patient.

---

Supplementary Table of Authors from the UCSF COMET Consortium

| Name | Institution |
| --- | --- |
| Cathy Cai | Department of Pathology and ImmunoX, UCSF, San Francisco, California, USA. |
| Jenny Zhan | Department of Pathology and ImmunoX, UCSF, San Francisco, California, USA. |
| Bushra Samad | Department of Pathology and ImmunoX, UCSF San Francisco, California, USA. |
| Kwok W. Im | Department of Pathology and ImmunoX, UCSF San Francisco, California, USA. |
| Nina K. Serwas | Department of Pathology and ImmunoX, UCSF San Francisco, California, USA. |
| Suzanna Chak | Division of Pulmonary and Critical Care Medicine, Department of Medicine, UCSF, San Francisco, California, USA. |
| Rajani Ghale | Division of Pulmonary and Critical Care Medicine, Department of Medicine, UCSF, San Francisco, California, USA. |
| Jeremy Giberson | Division of Pulmonary and Critical Care Medicine, Department of Medicine, Zuckerberg San Francisco General Hospital and Trauma Center, UCSF, San Francisco, California, USA. |
| Ana Gonzalez | Division of Pulmonary and Critical Care Medicine, Department of Medicine, Zuckerberg San Francisco General Hospital and Trauma Center, UCSF, San Francisco, California, USA. |
| Alejandra Jauregui | Division of Pulmonary and Critical Care Medicine, Department of Medicine, UCSF, San Francisco, California, USA. |
| Deanna Lee | Division of Pulmonary and Critical Care Medicine, Department of Medicine, Zuckerberg San Francisco General Hospital and Trauma Center, Cardiovascular Research Institute, UCSF, San Francisco, CA, USA. |
| Viet Nguyen | Division of Pulmonary and Critical Care Medicine, Department of Medicine, Zuckerberg San Francisco General Hospital and Trauma Center, Cardiovascular Research Institute, UCSF, San Francisco, CA, USA. |
| Kimberly Yee | Division of Pulmonary and Critical Care Medicine, Department of Medicine, University of California San Francisco, Cardiovascular Research Institute, UCSF, San Francisco, CA, USA. |
| Yumiko Abe-Jones | Division of Hospital Medicine, UCSF, San Francisco, California, USA. |
| Logan Pierce | Division of Hospital Medicine, UCSF, San Francisco, California, USA. |
| Priya Prasad | Division of Hospital Medicine, UCSF, San Francisco, California, USA. |
| Pratik Sinha | Division of Pulmonary and Critical Care Medicine, Department of Medicine, UCSF, San Francisco, California, USA. |
| Alexander Beagle | Department of Medicine, UCSF San Francisco, California, USA |
| Tasha Lea | Department of Pathology, UCSF San Francisco, California, USA. |
| Armond Esmalii | Division of Hospital Medicine, University of California, San Francisco, CA, USA. |
| Austin Sigman | Division of Pulmonary and Critical Care Medicine, Department of Medicine, University of California San Francisco, San Francisco, California, USA. |
| Gabriel M Ortiz | Department of Medicine, Zuckerberg San Francisco General Hospital and Trauma Center, University of California San Francisco |
| Kattie Raffel | Division of Hospital Medicine, University of California, San Francisco, CA, USA. |
| Chayse Jones | Division of Pulmonary and Critical Care Medicine, Department of Medicine, University of California San Francisco, San Francisco, California, USA. |
| Kathleen Liu | Division of Nephrology, Department of Medicine, University of California at San Francisco School of Medicine, San Francisco, CA, United States<br>Division of Critical Care Medicine, Department of Anesthesia, University of California at San Francisco School of Medicine, San Francisco, CA, United States. |
| Walter Eckalbar | Division of Pulmonary and Critical Care Medicine, Department of Medicine, Cardiovascular Research Institute and CoLabs, UCSF, San Francisco, CA, USA. |

## Notes

### Competing Interest Statement

The authors have declared no competing interest.

## References

1. D. Mathew et al., Deep immune profiling of COVID-19 patients reveals distinct immunotypes with therapeutic implications. Science 369, (2020).

2. J. Schulte-Schrepping et al., Severe COVID-19 Is Marked by a Dysregulated Myeloid Cell Compartment. Cell 182, 1419–1440 e1423 (2020).

3. J. Hadjadj et al., Impaired type I interferon activity and inflammatory responses in severe COVID-19 patients. Science 369, 718–724 (2020).

4. R. Zilionis et al., Single-Cell Transcriptomics of Human and Mouse Lung Cancers Reveals Conserved Myeloid Populations across Individuals and Species. Immunity 50, 1317–1334 e1310 (2019).

5. C. Trapnell et al., The dynamics and regulators of cell fate decisions are revealed by pseudotemporal ordering of single cells. Nat Biotechnol 32, 381–386 (2014).

6. I. C. Huang et al., Distinct patterns of IFITM-mediated restriction of filoviruses, SARS coronavirus, and influenza A virus. PLoS Pathog 7, e1001258 (2011).

7. M. Reyes et al., An immune-cell signature of bacterial sepsis. Nat Med 26, 333–340 (2020).

8. B. Engelmann, S. Massberg, Thrombosis as an intravascular effector of innate immunity. Nat Rev Immunol 13, 34–45 (2013).

9. S. Cui, S. Chen, X. Li, S. Liu, F. Wang, Prevalence of venous thromboembolism in patients with severe novel coronavirus pneumonia. J Thromb Haemost 18, 1421–1424 (2020).

10. P. Davizon-Castillo, J. W. Rowley, M. T. Rondina, Megakaryocyte and Platelet Transcriptomics for Discoveries in Human Health and Disease. Arterioscler Thromb Vasc Biol 40, 1432–1440 (2020).

11. K. D. Mason et al., Programmed anuclear cell death delimits platelet life span. Cell 128, 1173–1186 (2007).

12. D. Bongiovanni et al., Transcriptome Analysis of Reticulated Platelets Reveals a Prothrombotic Profile. Thrombosis and haemostasis 119, 1795–1806 (2019).

13. W. S. Chen et al., Uncovering axes of variation among single-cell cancer specimens. Nat Methods 17, 302–310 (2020).

14. E. Pujadas et al., SARS-CoV-2 viral load predicts COVID-19 mortality. Lancet Respir Med 8, e70 (2020).

15. S. Hue et al., Uncontrolled Innate and Impaired Adaptive Immune Responses in Patients with COVID-19 ARDS. Am J Respir Crit Care Med, (2020).

16. Y. Wang et al., Kinetics of viral load and antibody response in relation to COVID-19 severity. J Clin Invest 130, 5235–5244 (2020).

17. P. Bastard et al., Auto-antibodies against type I IFNs in patients with life-threatening COVID-19. Science, eabd4585 (2020).

18. S. Sammicheli et al., Inflammatory monocytes hinder antiviral B cell responses. Sci Immunol 1, (2016).

19. H. Huang, C. Benoist, D. Mathis, Rituximab specifically depletes short-lived autoreactive plasma cells in a mouse model of inflammatory arthritis. Proc Natl Acad Sci U S A 107, 4658–4663 (2010).

20. R. J. Looney, J. Huggins, Use of intravenous immunoglobulin G (IVIG). Best Pract Res Clin Haematol 19, 3–25 (2006).

## References

1. D. Bongiovanni et al., Transcriptome Analysis of Reticulated Platelets Reveals a Prothrombotic Profile. Thrombosis and haemostasis 119, 1795–1806 (2019).

2. T. Stuart et al., Comprehensive Integration of Single-Cell Data. Cell 177, 1888–1902 e1821 (2019).

3. D. Dominguez et al., A high-resolution transcriptome map of cell cycle reveals novel connections between periodic genes and cancer. Cell research 26, 946–962 (2016).

4. H. M. Kang et al., Multiplexed droplet single-cell RNA-sequencing using natural genetic variation. Nat Biotechnol 36, 89–94 (2018).

5. C. Genomes Project et al., A global reference for human genetic variation. Nature 526, 68–74 (2015).

6. H. Li, A statistical framework for SNP calling, mutation discovery, association mapping and population genetical parameter estimation from sequencing data. Bioinformatics 27, 2987–2993 (2011).

7. C. Hafemeister, R. Satija, Normalization and variance stabilization of single-cell RNA-seq data using regularized negative binomial regression. Genome biology 20, 296 (2019).

8. C. S. McGinnis, L. M. Murrow, Z. J. Gartner, DoubletFinder: Doublet Detection in Single-Cell RNA Sequencing Data Using Artificial Nearest Neighbors. Cell Syst 8, 329–337 e324 (2019).

9. I. Korsunsky et al., Fast, sensitive and accurate integration of single-cell data with Harmony. Nat Methods 16, 1289–1296 (2019).

10. X. Qiu et al., Reversed graph embedding resolves complex single-cell trajectories. Nat Methods 14, 979–982 (2017).

11. X. Qiu et al., Single-cell mRNA quantification and differential analysis with Census. Nat Methods 14, 309–315 (2017).

12. C. Trapnell et al., The dynamics and regulators of cell fate decisions are revealed by pseudotemporal ordering of single cells. Nat Biotechnol 32, 381–386 (2014).

14. K. R. Moon et al., Visualizing structure and transitions in high-dimensional biological data. Nat Biotechnol 37, 1482–1492 (2019).

15. C. R. Zamecnik et al., ReScan, a Multiplex Diagnostic Pipeline, Pans Human Sera for SARS-CoV-2 Antigens. medRxiv, 2020.2005.2011.20092528 (2020).

16. A. Leligdowicz et al., Validation of two multiplex platforms to quantify circulating markers of inflammation and endothelial injury in severe infection. PLoS One 12, e0175130 (2017).

